# Poising and connectivity of emergent human developmental enhancers in the transition from naive to primed pluripotency

**DOI:** 10.1101/2025.10.02.679816

**Authors:** Marina C. Nocente, Monica Della Rosa, Andrew A. Malcolm, George Lister, Isabella Savin, Helen Ray-Jones, Sarah Elderkin, Ruilin Tian, Simon Andrews, Adam Bendall, Claudia I. Semprich, Martin Kampmann, Valeriya Malysheva, Maria Rostovskaya, Peter Rugg-Gunn, Mikhail Spivakov

**Author notes:** These authors contributed equally (Marina C. Nocente, Monica Della Rosa). AAM - Centre for Cancer Research, The Institute of Genetics and Cancer, Edinburgh, EH4 2XU, UK., GL and IS - Department of Biochemistry, University of Oxford, Oxford, OX1 3QU, UK., HRJ - Department of Internal Medicine, Erasmus MC, Rotterdam, The Netherlands., RT - School of Medicine, Southern University of Science and Technology, Shenzhen, Guangdong Province, 518055, China., CIS - Cambridge Stem Cell Institute, Jeffrey Cheah Biomedical Centre, Cambridge Biomedical Campus, University of Cambridge, Cambridge, UK.

## Abstract

In primed human pluripotent stem cells (hPSCs) resembling post-implantation epiblast, numerous lineage-specific enhancers assume the poised chromatin state, co-marked by H3K4me1 and Polycomb-associated H3K27me3 histone modifications. In contrast, poised enhancers (PEs) are scarce in naive hPSCs that model pre-implantation epiblast. PEs form abundant chromosomal contacts with developmental genes, but when these contacts emerge, how their formation relates to enhancer poising and their functional significance remains incompletely understood. Here, we devise high-resolution, PE-targeted Capture Hi-C to generate a comprehensive atlas of PE chromosomal contacts in the time course of hPSC transition from the naive to primed state. We find that enhancer poising emerges early in the transition, while the contacts show diverse dynamics that is only partially coupled to poising. PROTAC-induced degradation of Polycomb Repressive Complex 2 (PRC2) early in the transition weakens PE connectivity, while inhibition of its H3K27 methyltransferase activity does not, suggesting a non-catalytic role of Polycomb in supporting PE contacts. Notably, PE contacts persist after developmental activation or ectopic CRISPRa targeting and can mediate long-range gene induction. Together, these findings reveal the temporal and mechanistic principles of PE connectivity, highlighting a potential role of PE contacts in establishing gene expression patterns in human development.

## INTRODUCTION

Enhancer elements play a key role in spatiotemporal gene control and can reside up to megabases away from their target genes, reaching them through 3D chromosomal contacts (Schoenfelder and Fraser 2019; Ray-Jones and Spivakov 2021). Aberrations in enhancer activity or enhancer-promoter communication, for example, due to genetic variation within enhancers or perturbations of topological domain boundaries, can lead to developmental defects and contribute to common and rare diseases (Lupiáñez et al. 2015; Gupta et al. 2017; Rickels and Shilatifard 2018; Zaugg et al. 2022).

Cell lineages are characterised by unique patterns of enhancer activity, whose emergence during development orchestrates lineage-specific gene expression (Heintzman et al. 2009; Heinz et al. 2015; Huang et al. 2016). Active enhancers are typically marked by H3K4me1 and H3K27ac histone modifications, associated with Trithorax/MLL and p300/CPB co-activators respectively, and contact their target gene promoters in 3D. In contrast, inactive enhancers often lack these marks and promoter contacts (Creyghton et al. 2010; Calo and Wysocka 2013; Wang et al. 2016; Javierre et al. 2016; Schoenfelder and Fraser 2019).

During early embryogenesis and in pluripotent stem cells, many enhancers pass through an intermediate ‘poised’ chromatin state before their lineage-specific activation or repression, in which they are co-marked by H3K4me1 and by H3K27me3 associated with Polycomb Repressive Complexes (PRCs) (Rada-Iglesias et al. 2011; Crispatzu et al. 2021). In addition, poised enhancers (PEs) often overlap orphan CpG islands and bind p300/CBP, although they lack the H3K27ac mark catalysed by these complexes (Pachano et al. 2021).

Similarly to active enhancers, PEs abundantly engage in 3D chromosomal contacts (Schoenfelder et al. 2015b; Freire-Pritchett et al. 2017; Crispatzu et al. 2021; Dimitrova et al. 2022), including with cognate ‘poised’ gene promoters characterised by H3K4me3 and H3K27me3 histone modifications, as well as RNA polymerase II pausing (Azuara et al. 2006; Bernstein et al. 2006; Stock et al. 2007). The chromosomal contacts of PEs, particularly those linking them with the promoters of their eventual target genes, may therefore provide critical clues into the developmental origins of lineage-specific transcriptional programmes. However, when these contacts are first established during early development, to what extent their formation is driven by the acquisition of the poised chromatin state at enhancers, and how they contribute to gene regulation, remains unclear.

In mouse, poised chromatin emerges in the pluripotent epiblast after formation of this lineage in the pre-implantation blastocyst and before germ layer specification during gastrulation (Dahl et al. 2016; Liu et al. 2016; Zheng et al. 2016; Du et al. 2020; Xiang et al. 2020; Xia and Xie 2020; Wilkinson et al. 2023). During this period, many poised loci also establish chromatin contacts (Xiang et al. 2020), although some contacts are already detectable at the pre-implantation stage (Crispatzu et al. 2021).

The fast development of the mouse epiblast (∼ 48 h) (Kojima et al. 2014), however, limits the ability to study the establishment of enhancer poising and connectivity in this system. Human embryos have considerably longer peri-implantation epiblast development and other notable interspecies differences (Boroviak et al. 2015; Nakamura et al. 2016; Coussement et al. 2025), but probing chromatin dynamics in them remains technically challenging. Human pluripotent stem cells (hPSCs), which represent the *in vitro* counterparts of the human embryonic epiblast, provide important models for studying these processes in human development. Alternative culture conditions can capture two distinct states of the pluripotent epiblast: ‘naive’ hPSCs, which resemble the pre-implantation epiblast, and ‘primed’ hPSCs (commonly referred to as ‘conventional hPSCs’), which correspond to the post-implantation, pre-gastrulation epiblast (Thomson et al. 1998; Takashima et al. 2014; Theunissen et al. 2014; Guo et al. 2016). We previously reported an experimental system for transitioning naive hPSCs towards the characteristics of the primed state (‘naive-to-primed transition’), which, over the course of two weeks, recapitulates the transcriptional and epigenetic dynamics of epiblast development (Rostovskaya et al. 2019; Agostinho de Sousa et al. 2023).

Naive hPSCs show reduced H3K27me3 levels at developmental genes, fewer poised loci and scarce long-range contacts at these regions (Theunissen et al. 2014; Chovanec et al. 2021; Zijlmans et al. 2022). In contrast, poised enhancers and promoters, and contacts between them, are abundant in primed hPSCs (Theunissen et al. 2014; Freire-Pritchett et al. 2017; Chovanec et al. 2021; Zijlmans et al. 2022). The naive-to-primed hPSC transition, therefore, offers a temporally-resolved view into the critical developmental window, during which emergent developmental enhancers first acquire the poised chromatin state and chromosomal contacts– processes that have not been systematically investigated in any organism (Xiong and Zhu 2025).

Many enhancer-promoter contacts evade detection by global chromosome conformation capture techniques such as Hi-C and Micro-C due to limitations on feasible coverage and resolution of these analyses imposed by sequencing library complexity, prompting the development of targeted enrichment approaches for these techniques (Schoenfelder and Fraser 2019; Kempfer and Pombo 2020). Capture Hi-C enables the enrichment of Hi-C material for the chromosomal contacts anchored at regions of interest, such as promoters or enhancers (typically in the tens of thousands), based on their sequence and irrespective of their chromatin state. This approach results in a ∼35-100-fold enrichment in the detection of targeted contacts compared with conventional Hi-C at equivalent sequencing depth (Schoenfelder et al. 2015a; Mifsud et al. 2015; Javierre et al. 2016; Schoenfelder et al. 2018).

Here, to address how PE chromosomal contacts emerge and are regulated during early human development, we adapted our recently developed low-input Capture Hi-C protocol (Ho et al. 2021; Malysheva et al. 2022) to enrich Hi-C material for the contacts of enhancers found in the poised state in primed hPSCs. We used this approach, termed Poised Enhancer-targeted Capture Hi-C (PECHi-C), to profile PE connectivity in a time course over the naive-to-primed transition of hPSCs globally and at high resolution, generating a comprehensive data resource and revealing diverse dynamics of enhancer contacts in relation to poising. We further show that PE connectivity is weakened by PROTAC-induced degradation of PRC2 early in the naive-to-primed hPSC transition, but not by inhibition of PRC2’s H3K27me3 catalytic activity, pointing to a non-catalytic role of Polycomb in stabilising PE contacts. Finally, we consider the fate of PE contacts after enhancer activation upon hPSC differentiation or ectopic CRISPR activation (CRISPRa). We find that subsets of PE contacts are maintained in these settings, suggesting their role in ‘priming’ target genes for subsequent developmental activation. Together, our study provides an extensive characterisation of the connectivity of emergent developmental enhancers during a critical time window of early human development, and reveals the temporal and mechanistic principles underpinning their chromosomal contacts, highlighting a potential role of PE contacts in establishing gene expression patterns in human development.

## RESULTS

### Establishment of a Poised Enhancer Capture Hi-C system

To profile PE contacts globally and at high resolution, we set out to devise a Poised Enhancer-targeted Capture Hi-C system (PECHi-C) to enrich Hi-C libraries for PE-anchored contacts using synthetic biotinylated RNAs (Figure 1A, Figure S1A). Sequence-based enrichment has enabled profiling the connectivity of these regions in a manner independent of their chromatin state, both before and after their acquisition of the poised state.

**Figure 1:**
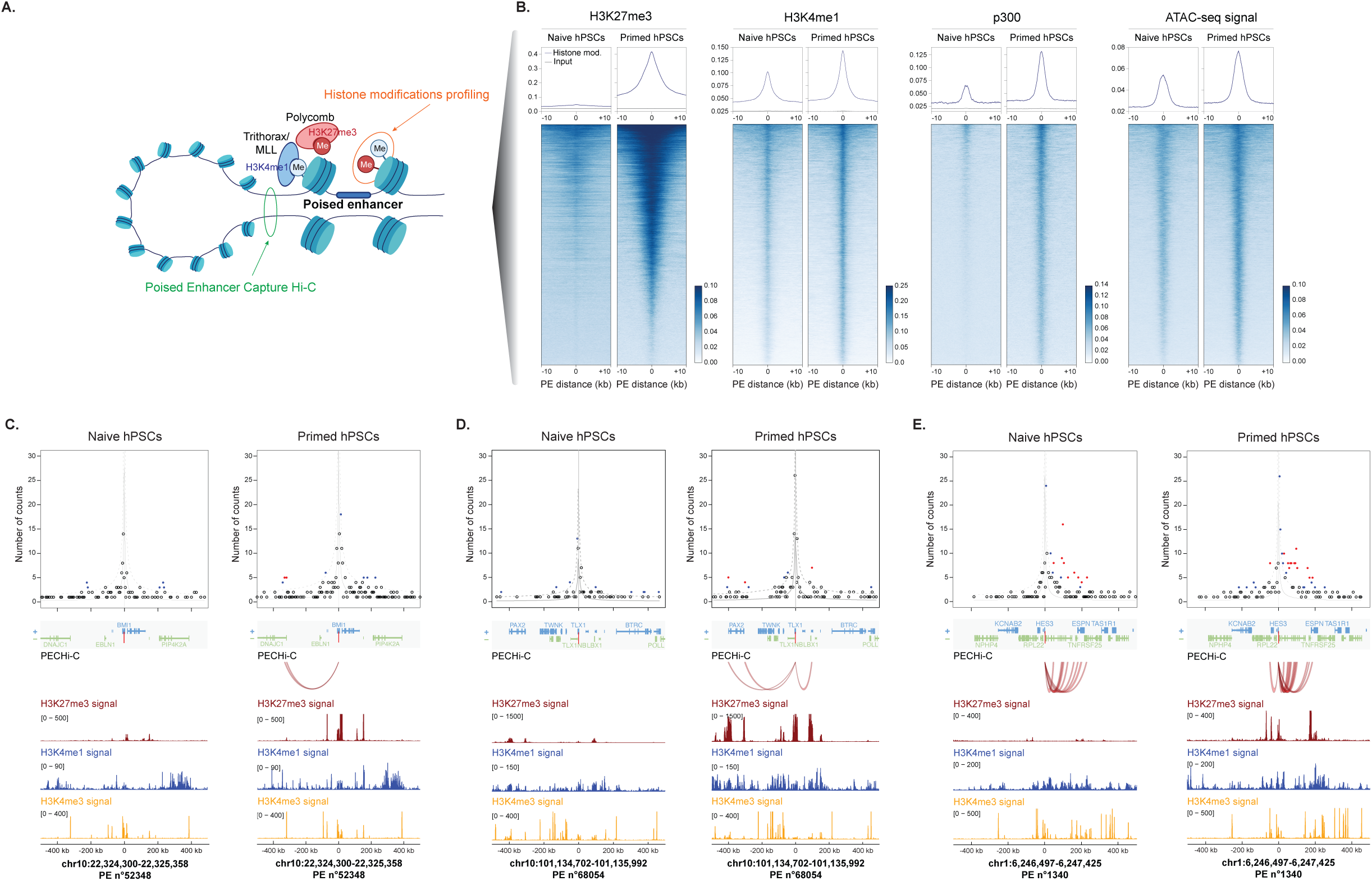
Identification of PEs and their chromosomal contacts in naive and primed hPSCs. (*A*) Schematic of a PE engaged in a chromosomal contact. PEs are characterised by the presence of H3K4me1 associated with Trithorax/MLL activators and H3K27me3 associated with Polycomb Repressive Complexes. This study used CUT&Tag for histone modifications to profile the acquisition of the poised state, and devised Poised Enhancer Capture Hi-C (PECHi-C) to profile PE chromosomal contacts. (*B*) Density plots (top) and heatmaps (bottom) showing averaged signal densities and signal enrichments per PE, respectively, for histone modifications (H3K27me3, H3K4me1), p300, and ATAC-seq (dark blue) versus input (grey) in naive and primed hPSCs. (*C, D, E*) Examples of chromosomal contacts detected for PE 52348 (*C*), 68054 (*D*), or 1340 (*E*) in naive and primed hPSCs. In (*C*) and (*D*), no significant contacts were detected in naive cells, while in (*E*), contacts were detectable in both naive and primed cells, with an overlap between both. Top panels: Normalised PECHiC read counts detected for each PE-harbouring baited *DpnII* fragment with ∼5kb bins of *DpnII* fragments with a 500 kb vicinity. Red dots indicate contacts with a CHiCAGO score ≥ 5, blue dots: contacts with a CHiCAGO score between 2 and 5, the rest of the fragment pairs are shown as black dots. Vertical gray lines indicate PE locations, with the mean background read counts and the upper boundaries of the 95% confidence intervals, estimated by CHiCAGO, indicated by solid and dotted gray lines, respectively. Gene annotations for the sense (+) and antisense (–) strands within the plotted regions are shown below, with red arcs indicating contacts with a CHiCAGO score ≥ 5. Bottom panels show CUT&Tag signals across the same genomic regions for H3K27me3 (dark red), H3K4me1 (dark blue), and H3K4me3 (orange).

Using chromatin profiling data from primed hPSCs, we identified a broad set of regions that capture all putative PE regions co-enriched for H3K4me1 and H3K27me3 and located more than 1 kb from the transcription start sites of protein-coding genes, and designed a sequence-capture system to target 32,142 *DpnII* fragments harbouring these loci (see Methods for details). While H3K4me1 signals were highly localised within PEs, H3K27me3 signals ranged from confined foci to broader domains of H3K27me3 enrichment (Figure 1B). Consistent with previous studies (Rada-Iglesias et al. 2011; Crispatzu et al. 2021; Pachano et al. 2021), most PEs were additionally bound by p300 and showed appreciable chromatin accessibility by ATAC-seq in primed hPSCs (Figure 1B). Also as expected from prior work (Theunissen et al. 2014; Chovanec et al. 2021; Zijlmans et al. 2022), in naive hPSCs, these elements were generally marked with H3K4me1, but had substantially reduced H3K27me3 and p300, as well as weaker ATAC-seq signals, compared to primed hPSCs (Figure 1B). Two thirds of these enhancers (66.8%) are in an active chromatin state in at least one human adult tissue according to data from the ENCODE project (Rozowsky et al. 2023) (Figure S1B), and a subset of them showed tissue-specific activity *in vivo* according to the VISTA Enhancer Browser (Visel et al. 2007; Kosicki et al. 2025) (Figure S1C). These observations support a model whereby many active enhancers in embryonic and adult tissues emerge through a poised state in the early stages of human development.

As a proof of concept, we initially applied PECHi-C in embryo-derived naive and primed hPSC lines. In naive hPSCs, we detected 48,038 high-confidence contacts (CHiCAGO score >= 5), linking 18,198 captured PEs with 43,839 genomic loci. In primed hPSCs, we identified 62,210 high-confidence contacts, linking 20,513 PEs with 54,163 loci. Examples of contacts detectable in both naive and primed cells or in primed cells alone are shown in Figure 1C, D, E. Notably, in primed hPSCs, the regions contacted by PEs showed enrichment for poising-associated histone marks H3K4me1/3 and H3K27me3 (Figure S1D). In contrast, in naive cells, where most of these enhancers have not yet acquired the poised state, their contacted regions were not enriched for H3K27me3 and only mildly enriched for H3K4me1 (Figure S1D).

Jointly, these results establish a Capture Hi-C system to study PE connectivity and suggest increased contacts of the targeted enhancers in primed hPSCs, where they are found in the poised state.

### The dynamics of enhancer poising and connectivity upon the naive-to-primed hPSCs transition

We next performed PECHi-C over seven time points during the transition of naive hPSCs towards the primed state (at days 1, 3, 5, 7, 10 and 14) using our previously published experimental system (Rostovskaya et al. 2019) (Figure 2A). Across these seven time points, we detected 323,112 unique significant contacts (CHiCAGO score >5 in at least one time point). As expected, the PE contact profiles at day 14 of naive-to-primed transition were the closest to, albeit still distinct from, those in conventional primed hPSCs (Figure S2A). For the remainder of the study, we focused on PE contacts with gene promoters and other PEs, as we reasoned that these contacts were most relevant for gaining insight into PE gene regulatory function. These included 8,477 contacts between 7,933 PEs and 5,794 promoters, and 1,626 contacts between pairs of captured PE regions.

**Figure 2:**
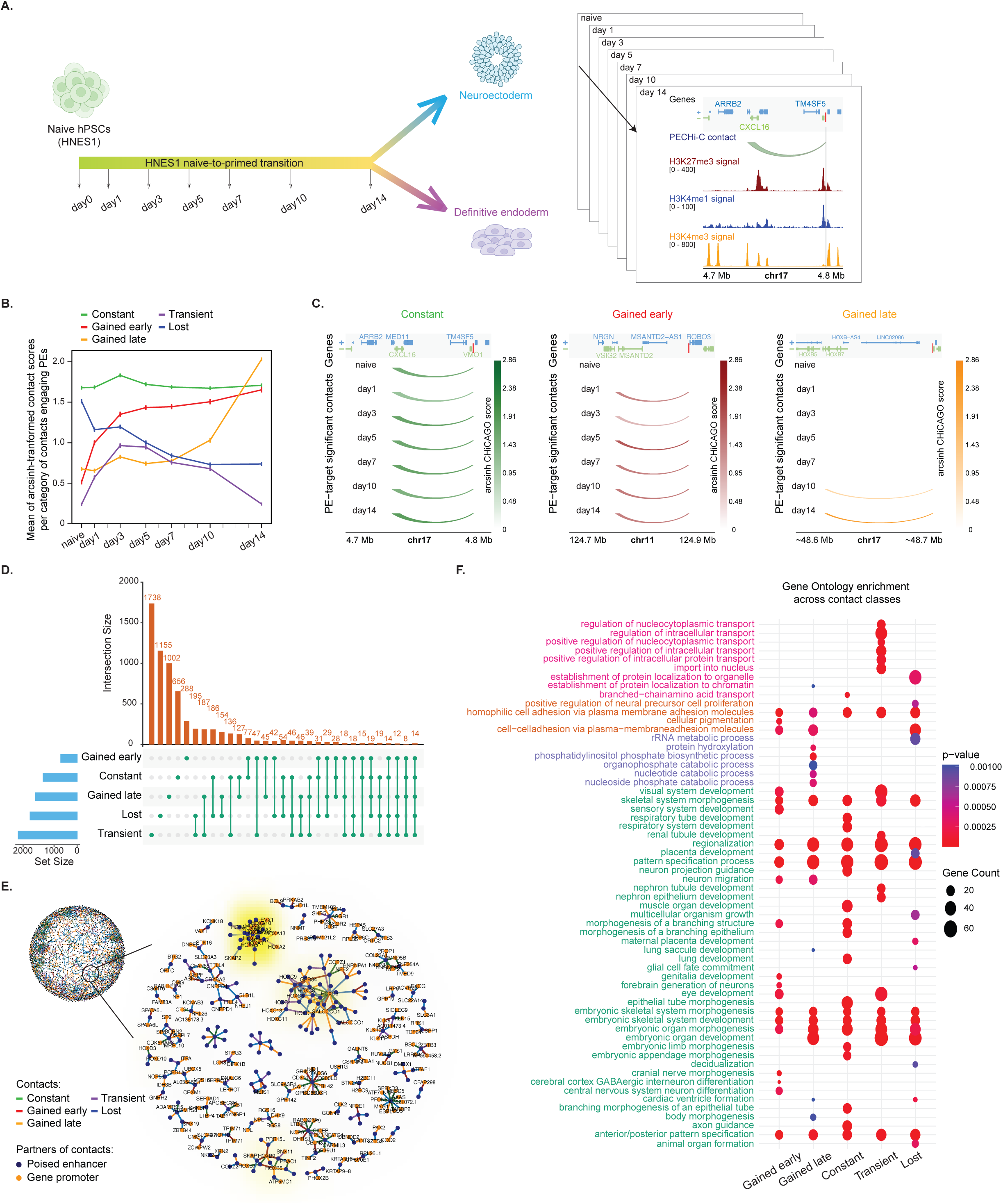
Dynamics of PE contacts during the naive-to-primed hPSCs transition. (*A*) Schematic of the transition from naive hPSCs to the primed state. Cells were collected at seven time points (naive = day 0, day 1, day 3, day 5, day 7, day 10, and day 14), as well as further differentiated into either neuroectoderm (NE) or definitive endoderm (DE) and analysed with CUT&Tag for histone modifications and PECHi-C (green arc) to detect PE chromosomal contacts. (*B*) Dynamics of PE chromosomal contacts with gene promoters and other PEs during the naive-to-primed transition. Mean arcsinh-tranformed CHiCAGO contact scores for each of the five temporal classes of PE contacts defined using Dynamic Time Warping clustering: ‘constant’ (green), ‘gained early’ (red), ‘gained late’ (orange), ‘transient’ (purple), and ‘lost’ (blue). Error bars represent standard deviations. (*C*) Representative examples of PE contacts during the naive-to-primed transition. Arcs represent PECHi-C contacts between a PE and a gene promoter within the indicated genomic region, with their principal colour reflecting temporal class annotation from panel B and colour intensities representing arcsinh-transformed CHiCAGO scores (see colour key). Absence of an arc corresponds to the CHiCAGO score of zero. Left: a ‘constant’ contact between PE 201999 and the *CXCL16* promoter. Middle: a ‘gained-early’ contact between PE 100396 and the *MSANTD2* promoter. Right: a ‘gained-late’ contact between PE 211333 and the *HOXB8* promoter. (*D*) UpSet plot showing the numbers of PEs involved in different classes of contact dynamics (orange bars, with green points indicating contact class combinations for each bar) and the total number of contacts per class (blue bars). (*E*) Visualisation of PE contact rewiring. Network plots showing the connectivity of PEs with rewired contacts (i.e., containing at least one ‘lost’ contact and one ‘gained early’ or ‘gained late’ contact). PEs are shown as blue nodes, gene promoters are shown in orange, and edges, colour-coded by class annotation from panel B, represent contacts with a CHiCAGO score >= 5 with gene promoters or other PEs. Top left: the total network of rewired PE contacts. Bottom right: a ‘zoomed-in’ plot focusing on 50 randomly sampled disjointed subnetworks, supplemented by the subnetworks involving the *HOX* loci, which are characterised by extensive connectivity and rewiring. The density of *HOX* genes in each subnetwork is colour-coded by yellow halos. (*F*) Gene Ontology enrichment analysis of genes contacted by PEs across different contact dynamics during the transition. Pathways are colour-coded by the broader terms curated based on these categories: molecular transport and localization (pink), cell adhesion and proliferation (orange), metabolic processes (blue), and development-related processes (green). Dot size represents gene counts, and dot colour reflects enrichment significance level (hypergeometric test p-value).

To assess the temporal dynamics of PE contacts upon the naive-to-primed transition, we clustered the contacts based on their CHiCAGO interaction scores using Dynamic Time Warping (DTW) (Alexis Sarda-Espinosa 2015). DTW is a distance metric developed to better capture similarities in temporal patterns between observations (in our case, the PEs’ contact scores) compared with standard metrics such as Euclidean distance. This analysis revealed five diverse classes of contacts: the largely timepoint-invariant ‘constant’ contacts (n = 1,999); contacts that were gained either early or late in the naive-to-primed transition (n = 781 and n = 2,045, respectively); transient contacts that peaked around day 3 (n = 2,847); and finally, contacts that were gradually lost in the course of the transition (n = 2,326) (Figures 2B, C, Figure S2B; Table S1).

While over two-thirds of PEs (4,452/6,445, 69%) contacted only a single gene promoter or PE, 80% of the remaining PEs (1,594/1,993) engaged in combinations of contacts from different temporal classes, indicating dynamic contact rewiring (Figure 2D). For example, 187 PEs formed both ‘gained-late’ and ‘transient’ contacts, 77 PEs both ‘gained-early’ and ‘constant’ contacts, and 42 PEs both ‘gained-early’ and ‘lost’ contacts. Finally, 14 PEs engaged in all five classes of contacts (Figure 2D). Network analysis highlighted highly-connected hubs of rewired PE contacts, including, in particular, with the promoters of *HOX* genes known to undergo extensive 3D remodelling during development (Figure 2E) (Noordermeer et al. 2014).

We asked whether genes in different temporal classes of PEs contacts had distinct functional annotations. Generally, all classes of contacts linked PEs with developmental genes (Figure 2F). However, the ‘gained-late’ contacts were additionally enriched for metabolic genes, while ‘transient’ contacts showed an enrichment for nuclear transport genes (Figure 2F), highlighting potentially distinct functional roles of PE contacts with different temporal dynamics.

In conclusion, this analysis has revealed diverse dynamics of PE contacts, including abundant acquisition and rewiring of contacts with developmental gene promoters upon the naive-to-primed transition.

### Chromatin features associated with poised enhancer contact dynamics

We next asked how the dynamics of enhancer connectivity relate to their acquisition of the poised state. To this end, we used concurrently generated CUT&Tag data for the poising-associated H3K27me3 and H3K4me1 across the same time course as our PECHiC samples (M.D.R., M.R., et al., in preparation) (Figures 2A, 3A, B).

**Figure 3:**
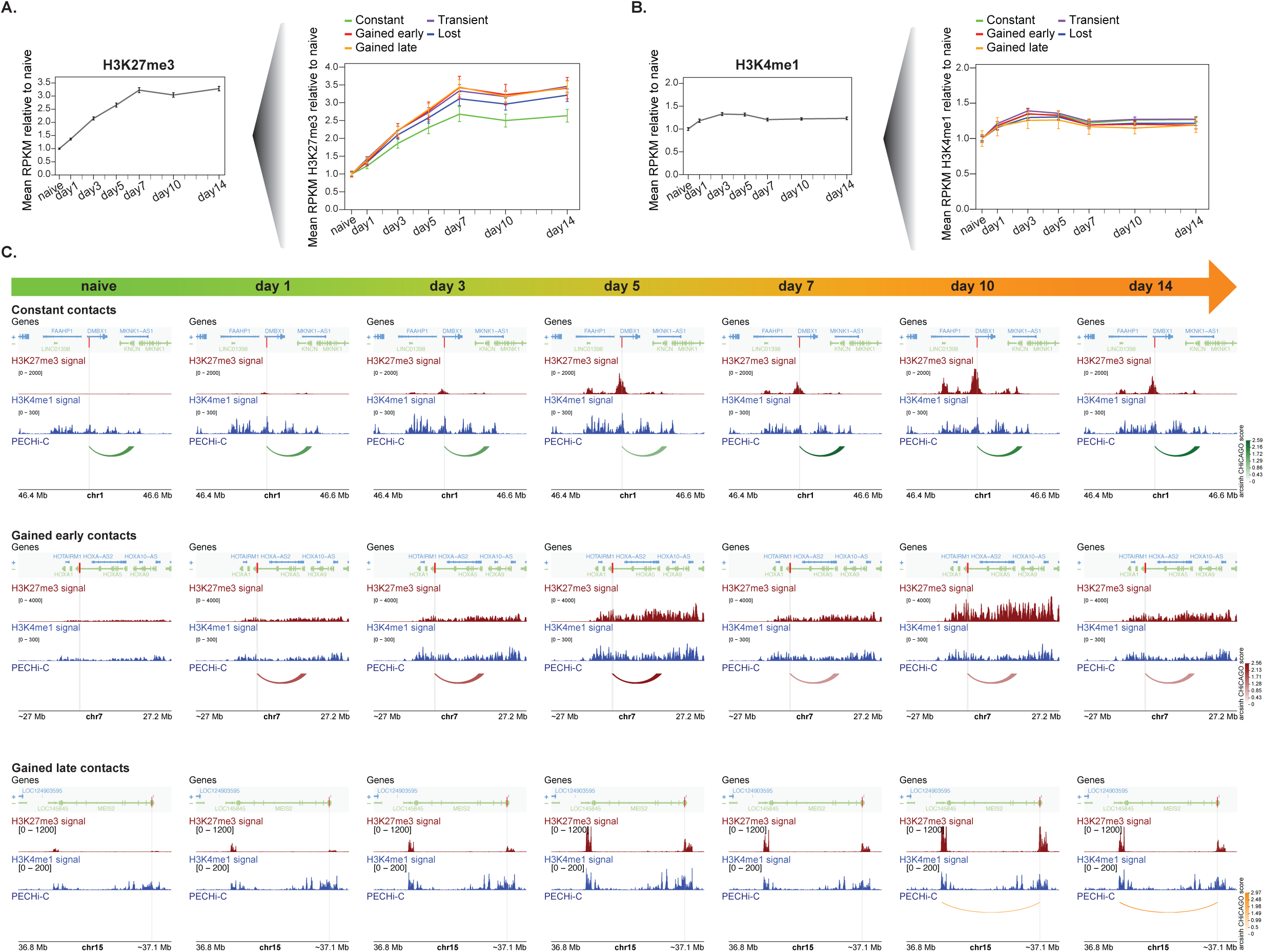
Dynamics of enhancer poising during the naive-to-primed transition. (*A, B*). The dynamics of the poising-associated histone modifications H3K27me3 (*A*) and H3K4me1 (*B*) during the naive-to-primed transition profiled by CUT&Tag. Left panels: Mean RPKM-normalized signals across PEs per time point, normalised to the levels detected in the naive state. Right panels: The same values averaged for PEs engaged in each temporal class of contacts. Error bars represent standard deviations. (*C*) Representative examples of CUT&Tag signals for H3K27me3 (dark red) and H3K4me1 (dark blue) at candidate PEs engaged in different classes of chromosomal contacts (‘constant’: top, ‘gained early’: middle, ‘gained late’: bottom) during the naive-to-primed transition. Arcs represent PECHi-C contacts between a PE and a gene promoter or another PE within the indicated genomic region. The arcs’ principal colour reflects temporal class annotation from Figure 2B and their colour intensities represent arcsinh-transformed CHiCAGO scores (see colour key). Absence of an arc corresponds to the CHiCAGO score of zero. Red and grey bars highlight the locations of PE baits.

In contrast to the highly diverse dynamics of PE contacts, we observed a consistent increase in H3K27me3 enrichment at the emergent PEs during the transition, which reached, by day 7, a plateau at ∼3.5-fold the levels observed in naive cells (Figure 3A, left panel). In contrast, H3K4me1 levels at these elements remained generally invariant throughout the transition (Figure 3B).

We found appreciable differences in H3K27me3 dynamics between PEs engaged in different temporal classes of contacts. In particular, PEs forming the ‘gained-early’ and ‘gained-late’ contacts generally showed the steepest increase in H3K27me3 during the transition (Figures 3A, right panel; 3C), while those engaged in constant contacts showed the smallest increase in H3K27me3 (Figures 3A, C). Overall, however, differences in H3K27me3 dynamics between subsets of PE could not fully account for their different contact dynamics.

To further investigate PE properties associated with the differential dynamics of their contacts, we assessed the breadth of H3K27me3 signals around PEs in the primed state (“H3K27me3 spreading”), as well as the identity of chromatin proteins recruited to these regions. To this end, we curated and uniformly processed public data for the architectural protein CTCF (Ji et al. 2016), pluripotency-associated transcription factors (TFs) OCT4, NANOG, SOX2, and TFAP2C (Ji et al. 2016; Chovanec et al. 2021; Osnato et al. 2021; Pastor et al. 2018; Dong et al. 2020), and co-factors Mediator and SWI/SNF (subunits BRM1, BRG1 and BAF155) (Ji et al. 2016; Huang et al. 2021) in naive and primed hPSCs. Finally, we used recently generated in-house data for the Polycomb-associated DPPA2 and DPPA4 transcription factors (Eckersley-Maslin et al. 2020; Gretarsson and Hackett 2020), which may facilitate Polycomb recruitment to chromatin (Malcolm et al., in revision).

Partitioning the captured PEs based on these properties revealed five coherent clusters (Figure 4A; Table S2). PEs in the first cluster mapped within the broadest areas of H3K27me3 enrichment and showed strong binding of Polycomb-associated DPPA2/4, as well as moderate occupancy of CTCF, pluripotency-associated TFs and co-factors (‘Broad H3K27me3’, n = 6,871; Figures 4A, B, Figure S2C). In contrast, PEs in the ‘CTCF + TFs’ cluster (n = 5,828) recruited CTCF and pluripotency TFs, but had tightly localised H3K27me3 signals and low DPPA2/4 occupancy (Figures 4A, B, Figure S2C). ‘Intermediate’ PEs (n = 9,524) fell within medium-width H3K27me3 signals, had moderate occupancy of DPPA2/4 and low levels of CTCF and pluripotency TFs. Finally, two clusters (n = 4,072 and n = 5,764 PEs) were characterised by high and low occupancy of most profiled factors, respectively (Figures 4A, B, Figure S2C). Notably, PEs in the ‘Broad H3K27me3’ and ‘High Occupancy’ clusters were particularly enriched for orphan CpG islands (Figure S2D).

**Figure 4:**
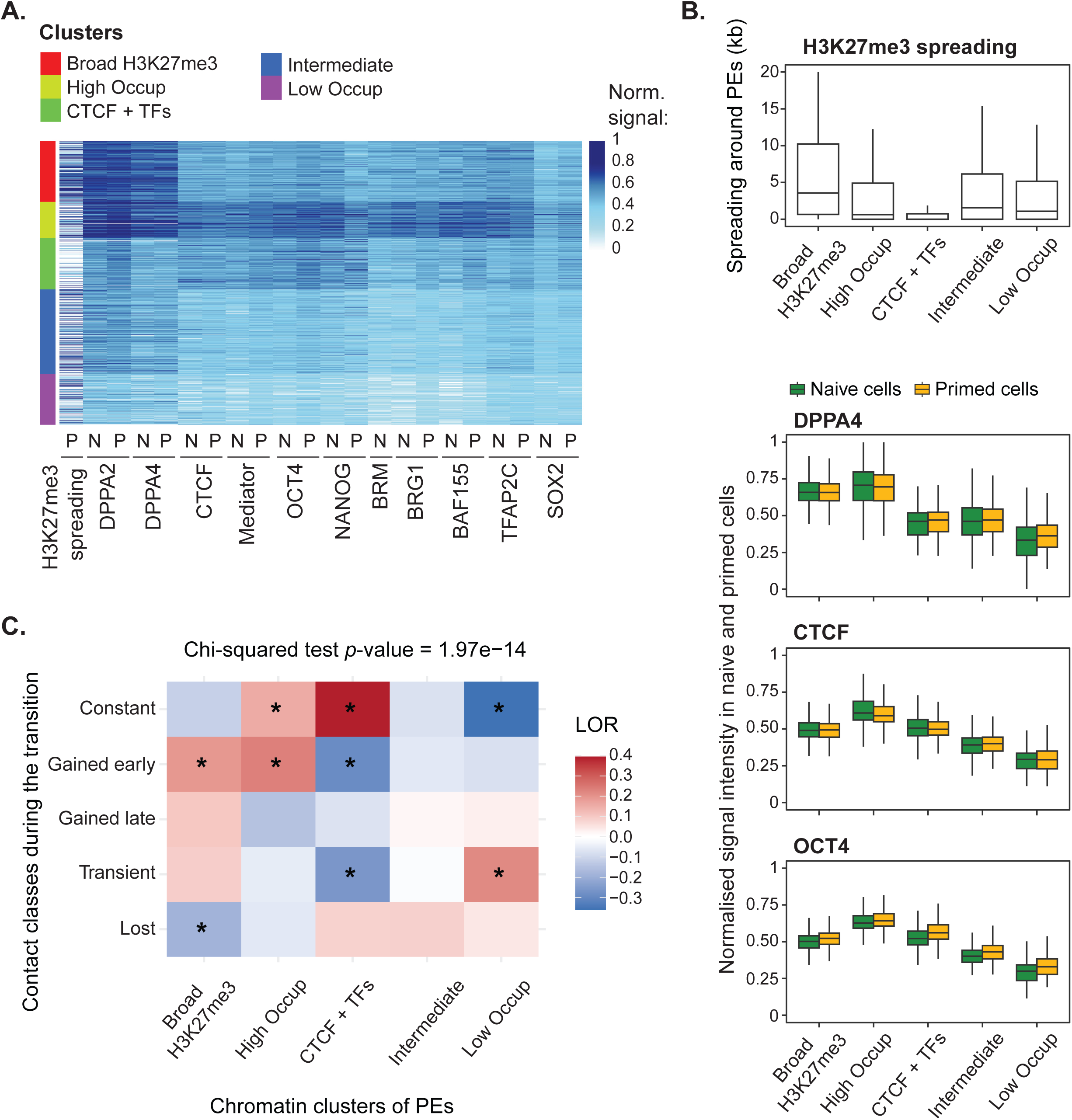
Association between PE chromatin features and their contact dynamics during the naive-to-primed transition. (*A*) Heatmap showing the extent of H3K27me3 spreading around PEs in primed cells, as well as the enrichment signals of transcription factors (DPPA2, DPPA4, CTCF, OCT4, NANOG, SOX2, TFAP2C) and cofactors (mediator, BRM, BRG1, BAF155) at all analysed PEs (rows) in naive and primed hPSCs. All values shown are normalised to the top 0.01 percentile for each dataset. Vertical labels denote PE partitioning into five ‘chromatin’ clusters based on these data: the “Broad H3K27me3” cluster (red, *n* = 6,871), the “High Occupancy” cluster (light green, *n* = 4,072), the “CTCF + TFs” cluster (green, n = 5,828), the “Intermediate” cluster (blue, *n* = 9,524), and the “Low Occupancy” cluster (purple, *n* = 5,764). (*B*) *Top:* The extent of H3K27me3 spreading around the PEs assigned to different chromatin clusters in primed cells. *Bottom:* Normalised signal intensities of DPPA4, CTCF, and OCT4 across PE chromatin clusters in naive (green) and primed (yellow) cells. Note that as these data were used in cluster definition, conventional statistical testing for their differences between clusters is not appropriate due to “double-dipping”. (*C*) Heatmap showing the enrichment (or depletion) of PE chromatin clusters (x-axis) for each class of temporal contact dynamics (y-axis). Colours represent log-odds ratios (LOR), stars indicate absolute chi-square standardised residuals above 1.96 reflecting significant enrichment or depletion.

PEs in these five clusters showed significant differences in the dynamics of chromosomal contacts (Figure 4C, Chi-squared test *p*-value = 1.97e-14). In particular the ‘Broad H3K27me3’ and ‘High Occupancy’ PEs were most enriched for the ‘gained-early’ contacts that emerged concomitantly with enhancer poising (Figure 4C). Consistent with this, PEs engaged in the ‘gained-early’ contacts tended to localise within the broadest areas of H3K27me3 enrichment (Supplementary Figure S2E). In contrast, the ‘constant’ contacts were strongly overrepresented in the ‘CTCF + TFs’ PE cluster, potentially implicating CTCF/cohesin-mediated chromatin loop extrusion in supporting these contacts. Notably, this cluster was also enriched for contacts that were lost upon the acquisition of poising. Finally, ‘low occupancy’ PEs were enriched for ‘transient’ contacts, suggesting they could be underpinned by transient binding events by these or other factors during the naive-to-primed transition that are not captured by this analysis (Figure 4C).

Taken together, these results suggest that PE contact dynamics may be influenced by a functional interplay of molecular mechanisms implicating Polycomb and loop extrusion.

### PRC2 degradation, but not inhibition of H3K27me3 catalytic activity, weakens poised enhancer connectivity

To directly test the role of Polycomb in facilitating PE connectivity early in the naive-to-primed transition in genetically unperturbed cells, we took advantage of potent small-molecule tools that enable PRC2 perturbation: the UNC6852 PROTAC that induces the proteasomal degradation of the key PRC2 components EED, EZH2, and SUZ12 (‘PRC2 PROTAC’) and the UNC1999 catalytic inhibitor of the H3K27 methyltransferases EZH1/2 (‘PRC2 inhibitor’) (Xu et al. 2015; Potjewyd et al. 2020) (Figure 5A). Treatment concentrations for each compound were optimised to achieve substantial depletion of total H3K27me3 levels over five days of culture (from naive to day 5), without major effects on cell morphology or viability, in line with prior studies (Agostinho de Sousa et al. 2023) (Figure S3A, B).

**Figure 5:**
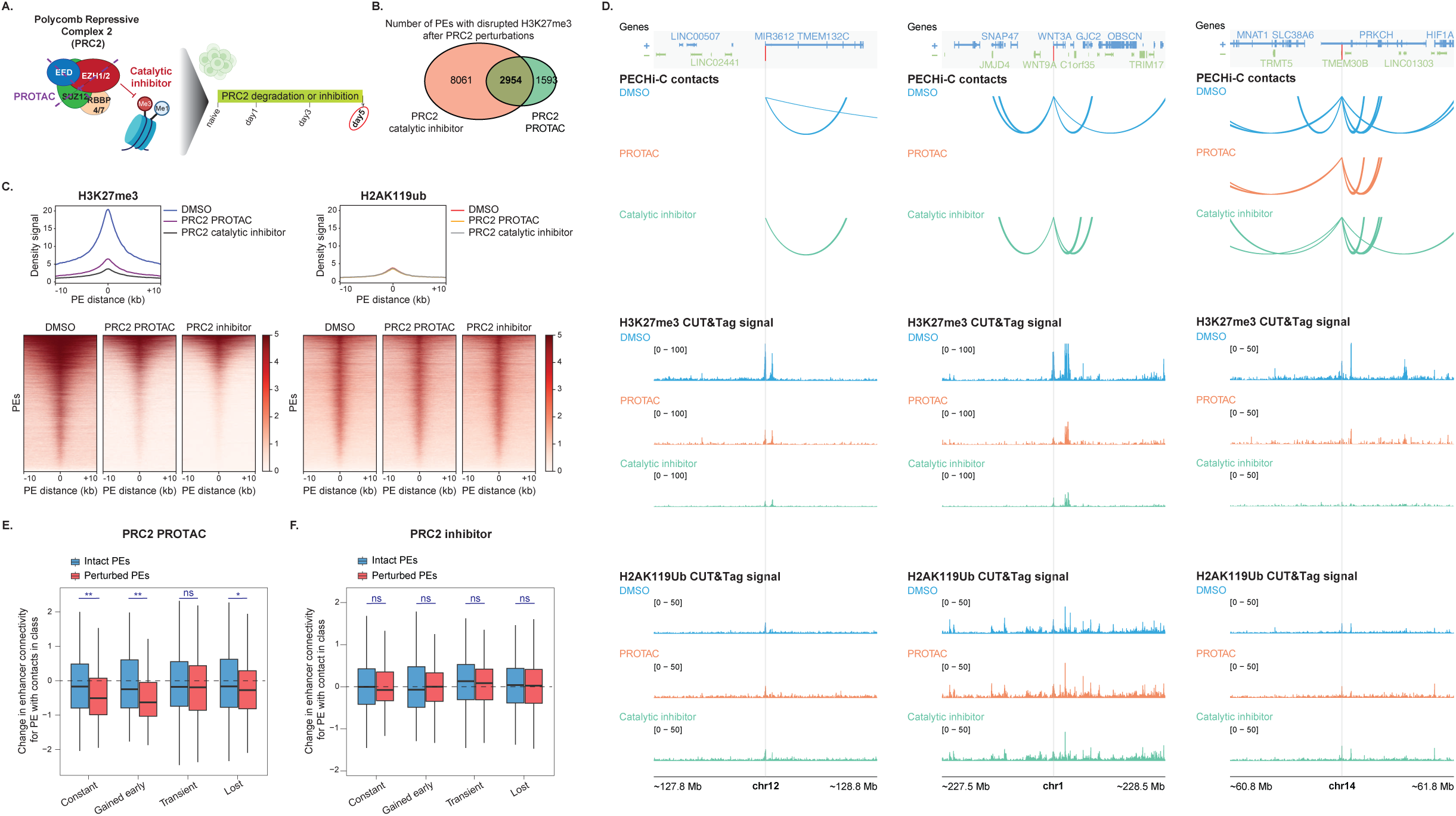
The effect of PRC2 perturbations on PE connectivity at day 5 of the naive-to-primed transition. (*A*) Schematic of the composition of Polycomb Repressive Complex 2 (PRC2) and the targets of utilised perturbations: a PRC2 PROTAC degrader (UNC6852) and a catalytic inhibitor of its EZH1/2 H3K27 methyltransferases (UNC1999). These compounds were added at the beginning of the transition and until day 5, when cells were harvested for analysis. (*B*) Venn diagram showing the numbers of PEs with the most robustly detected loss of H3K27me3 (‘perturbed PEs’) following PRC2 perturbations with either or both compounds compared to DMSO controls. (*C*) Density plots (top) and heatmaps (bottom) showing H3K27me3 and H2AK119ub signals across PE-centred 10-kb windows detected at day 5 of the transition following treatment with DMSO, PRC2 PROTAC, or PRC2 catalytic inhibitor from day 0. (*D*) Examples of loss of PECHi-C contacts following treatment with a PRC2 PROTAC (orange) or a PRC2 catalytic inhibitor (green), compared to DMSO (blue). Top panel: Gene locations, with arcs showing PE contacts in each condition. Middle and bottom panels: H3K27me3 (middle) and H2AK119ub (bottom) CUT&Tag signals in each condition. Grey and red bars highlight the locations of PE baits. (*E, F*) Changes in the connectivity (t-values) of PEs engaged in promoter/PE contacts of each temporal class after treatment with PRC2 PROTAC (*E*) or PRC2 catalytic inhibitor (*F*), for intact (blue) or perturbed (red) PEs. (‘Gained-late’ contacts were still weak at day 5 and not included in this analysis). The significance from Wilcoxon rank-sum tests is indicated as: ns = p-value > 0.05; * = 0.01 < p-value ≤ 0.05; ** = 0.001 < p-value ≤ 0.01).

We tested the effects of PRC2 PROTAC or PRC2 inhibitor treatment on PE chromatin state using calibrated CUT&Tag at day 5 of the naive-to-primed transition, at which H3K27me3 signals at PEs began to plateau in unperturbed conditions (Figure 3A). Both treatments significantly reduced H3K27me3 levels at large subsets of PEs (referred to as ‘perturbed PEs’ hereinafter) relative to DMSO. The most robust differences were observed at 4,547 PEs for PRC2 PROTAC and 11,015 for PRC2 inhibitor, with a shared subset of 2,954 perturbed PEs (DESeq2 padj<0.05; Figures 5B, C, D). In contrast, the levels of the PRC1-associated histone mark H2AK119ub remained largely unaffected after both treatments, suggesting that PRC1 is not affected by the degradation or catalytic inhibition of PRC2 at this time point (Figures 5C, D). Notably, the shared subset of perturbed PEs generally mapped to regions of broad H3K27me3 enrichment (Figure S3C). As expected, both treatments also induced changes in gene expression, with the PRC2 inhibitor resulting in expression changes for a larger subset of genes compared with the PROTAC (Figures S3D, E). However, since gene expression changes may result from either the effects of these perturbations on PEs or their direct effects on gene promoters, we did not consider these data in further analyses.

To assess how PRC2 perturbations affected PE connectivity, we performed PECHi-C in cells treated with PRC2 PROTAC, PRC2 inhibitor or DMSO and harvested at day 5 of the transition (Figure 5D and Figure S3F, top panels; Table S3). For a global analysis, we quantified changes in total PE connectivity upon PRC2 degradation or inhibition relative to DMSO treatment. We first assessed treatment-induced changes in the connectivity of the shared subset of 2,954 perturbed PEs and then compared them with those observed for the ‘intact PEs’ that were not significantly affected by either treatment. This way, intact PEs served as within-sample internal controls to ensure that any observed differences in the connectivity of perturbed PEs between treated and untreated cells could not be fully accounted for by batch effects.

PRC2 PROTAC treatment at day 5 led to an overall reduction in the connectivity of perturbed PEs compared with DMSO-treated controls (median *t-*value = -0.25; single-sample Wilcoxon test against *t =* 0, *p*-value = 7e-11; Figure S3G). While this effect was relatively mild, it was significantly more pronounced than for intact PEs (two-sample Wilcoxon test, *p*-value = 0.001; Figure S3G), indicative of a specific effect on enhancers losing the poised state upon PRC2 degradation. Furthermore, the magnitude of degradation-induced changes in connectivity varied among perturbed PEs engaged in distinct temporal classes of contacts (Figures 5E and S3H). Specifically, perturbed PEs engaged in ‘gained-early’ and ‘constant’ contacts showed the strongest reduction in connectivity upon PROTAC treatment relative to DMSO, whereas those engaged in ‘transient’ and ‘lost’ contacts exhibited milder changes (Figures 5E and S3H; ‘gained-late’ contacts were still weak at day 5 of transition and were not considered in this analysis). Expectedly, the connectivity of intact PEs upon PROTAC treatment also changed only mildly relative to DMSO and, for all classes but ‘transient’, significantly less than that of the respective perturbed PEs (Figure 5E).

In contrast to PRC2 depletion, inhibition of PRC2 catalytic activity had little impact on the overall connectivity of perturbed PEs relative to DMSO-treated controls, both overall (median *t-*value = 0.01; single-sample Wilcoxon test against *t =* 0, *p*-value = 0.197; Figure S3G) and across PEs engaged in different temporal classes of contacts (Figure 5F). There were also no significant differences between the effect of PRC2 inhibition on perturbed PEs relative to intact PEs (two-sample Wilcoxon test *p-*values>0.05; Figure 5F and Figure S3G).

Jointly, these results demonstrate that PRC2 supports the establishment of PE connectivity early in the naive-to-primed transition, potentially acting through mechanisms independent of its H3K27me3 catalytic activity.

### PE contacts gained before or during poising are retained after developmental enhancer activation

We next considered the function of PE contacts showing different temporal dynamics. Since PEs typically contact transcriptionally inactive genes, we asked whether their contacts were retained when these enhancers transitioned from the poised to an active state upon hPSC differentiation and could thus contribute to developmental gene induction. To assess this, we induced differentiation towards either definitive endoderm (DE) or neuroectoderm (NE) following naive-to-primed transition (Figure 2A). We then profiled enhancer contacts by PECHi-C, detecting 2,003 and 2,211 PE contacts with gene promoters or other PEs in DE and NE cells, respectively. We integrated these data with in-house chromatin profiles of DE and NE cells generated by CUT&Tag against H3K27me3, H3K4me1, H3K4me3 and H3K27ac (M. D. R., M. R., et al., in preparation).

The majority of PEs remained poised or became inactive (i.e., lost H3K4me1) upon differentiation to either embryonic tissue (Figure S4A, B), consistent with the DE and NE cells still representing early lineage progenitors. However, we focused on the 307 and 99 connected PEs that exchanged H3K27me3 for H3K27ac upon differentiation in DE and NE, respectively, reflecting their developmental activation (Figure 6A, Figure S4A, B and examples in Figure S4C, D). Notably, a significant proportion of promoters and enhancers contacted by these activated elements also lost H3K27me3 (56% and 37%, respectively, in DE and NE), and/or gained H3K4 mono- or trimethylation (39% and 55%, respectively), suggesting coordinated chromatin dynamics on both ends of the contacts.

**Figure 6:**
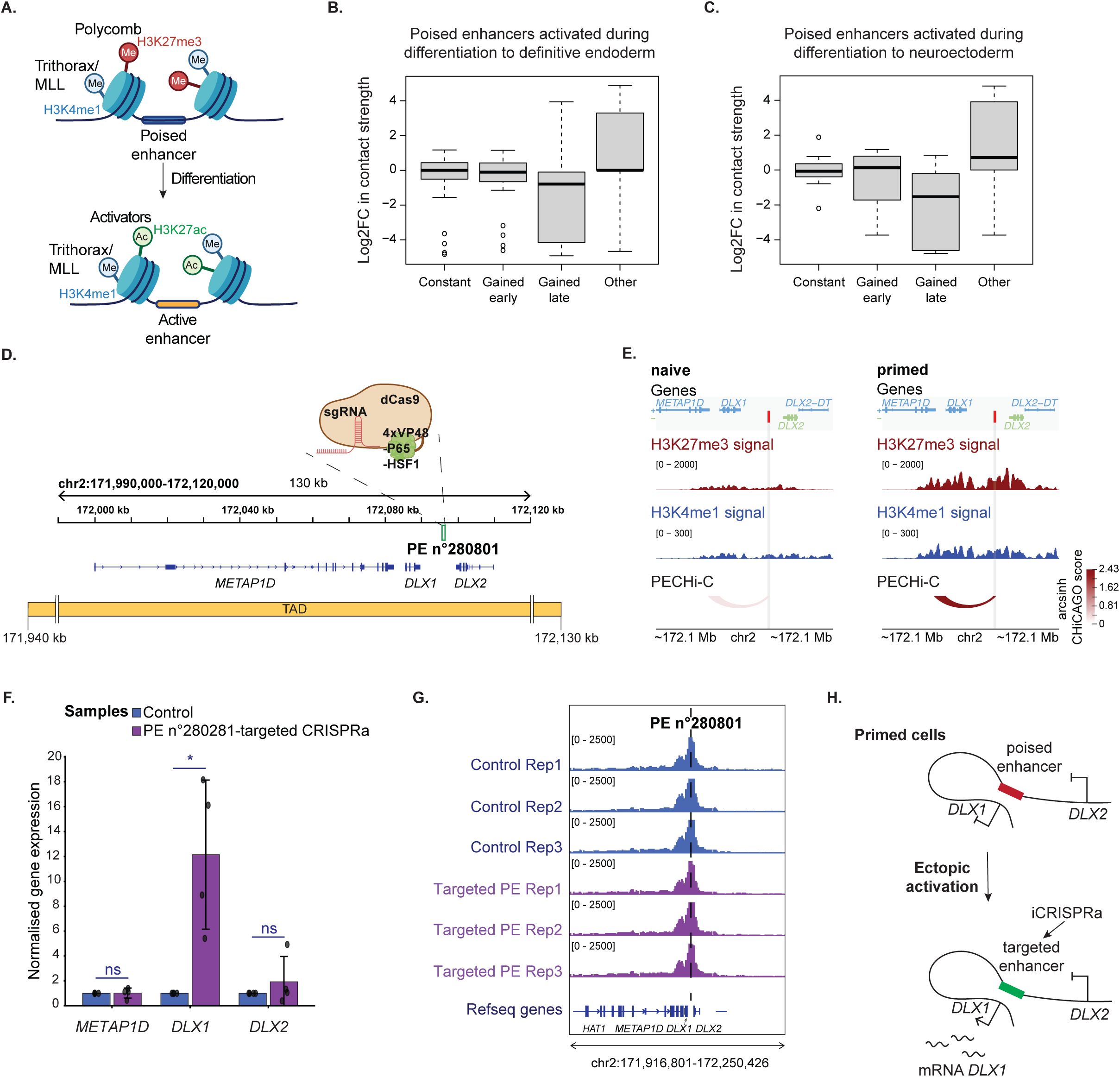
PE contacts persist after developmental activation or ectopic CRISPRa targeting and can mediate long-range gene induction. (*A*) Schematic of chromatin state transition upon PE activation, with the loss of the repressive H3K27me3 mark deposited by Polycomb and acquisition of the active H3K27ac mark deposited by histone acetyltransferases such as p300/CPB, while the H3K4me1 mark associated with Trithorax/MLL is retained. (*B, C*) Log2 fold change (Log2FC) in contact strength (proxied by arcsinh-CHiCAGO scores) for PEs involved in different types of contact dynamics that became active during differentiation to definitive endoderm (*B*) or neuroectoderm (*C*). (*D*) IGV screenshot of the *DLX1*-*DLX2* genomic region showing the position of PE 280801 (green rectangle) targeted by an inducible CRISPR activation (CRISPRa) system containing 4xVP48-P65-HSF1, dCas9, and a cocktail of single guide RNAs (sgRNA) targeting the poised enhancer 280801 in primed WTC11 hPSCs. The topological associated domain (TAD) encompassing the *DLX1*-*DLX2* genomic region is represented as an orange rectangle. (*E*) Visualisation of the H3K27me3 (dark red) and H3K4me1 (dark blue) CUT&Tag signals and the ‘gained-early’ PECHi-C contact (arcs) between PE 280801 (red bar) and the *DLX1* promoter in naive and primed cells, with arc color intensity reflecting arcsinh-transformed CHiCAGO scores. Red and grey bars highlight the locations of PE baits. (*F*) Normalised gene expression levels of *DLX1*, *DLX2* and *METAP1D* following inducible CRISPR activation (CRISPRa) of PE 280801 (purple) versus controls using scrambled gRNAs (blue. Error bars represent standard deviations (n = 4). (*G*) IGV visualisation of PE 280801 contacts detected at high resolution by 4C-seq following inducible CRISPR activation (purple) versus controls using scrambled gRNAs (blue). (*H*) Schematic of the effects of CRISPRa targeting to PE 280801 on the chromatin conformation and gene expression in the *DLX1*-*DLX2* locus, with the pre-existing PE-*DLX1* contact mediating *DLX1* activation, while the uncontacted *DLX2* remains unaffected.

We next assessed changes in contact strength following PE activation in DE and NE cells, stratifying these changes by the preceding temporal dynamics of the same contacts in the naive-to-primed transition (Table S4). We found that the dynamics of PE contacts during the transition significantly affected their retention upon enhancer activation (Kruskal–Wallis test *p*-values < 2.2e-16 in DE, and 1.42e-07 in NE). In particular, the contacts showing the ‘constant’ or ‘gained-early’ dynamics during the transition were largely retained by activated enhancers in DE, while ‘gained-late’ contacts were generally lost upon PE activation (Figure 6B and Figure S4C). A similar dynamic was seen in NE cells (Figure 6C and Figure S4D).

Therefore, PE contacts formed before or concomitantly with poising were retained upon the subsequent activation of these PEs during development, potentially contributing to gene induction.

### PE contacts mediate selective gene induction in the *DLX1/2* locus

To further probe whether PE contacts support gene induction, we targeted a CRISPR activation system (CRISPRa) to a PE connected to a developmental gene in primed hPSCs and assessed the effects of this perturbation on gene expression and PE connectivity.

For this analysis, we took advantage of WTC11 primed hPSCs containing a stably integrated, inducible CRISPRa system, including the dCas9 protein fused with a VPH activator (4 copies of VP48 fused with P65-HSF1) and a DHFR degron (Tian et al. 2021) (Figure 6D). Using this system, we targeted a PE (PECHi-C ID: 280801) located between two genes encoding neural-associated transcription factors *DLX1* and *DLX2* (∼26 kb and ∼12 kb away from their promoters, respectively), contained within the same topological associated domain (TAD) (Figures 6D, E). Importantly, while PE 280801 is connected to the *DLX1* promoter through a ‘gained-early’ contact, it is not detectably connected to the more proximal *DLX2* promoter (Figures 6E and S4E).

Targeting CRISPRa to PE 280801 activated the expression of the connected *DLX1* gene by >12-fold, but did not significantly affect the expression of *DLX2* or a more distal gene, *METAP1D*, located ∼36.7 kb upstream of *DLX1* (Figure 6F). To assess whether CRISPRa targeting resulted in changes in the connectivity pattern of PE 280801, we used *DpnII-*based 4C-seq with this PE as the viewpoint. This analysis confirmed the asymmetric pattern of PE 280801 connectivity under unperturbed conditions and showed that it was unaffected by CRISPR activation (Figure 6G).

Taken together, these results indicate that the contact between PE 280801 and *DLX1* can mediate this gene’s induction, while the unconnected, more proximal gene *DLX2* remains unaffected (Figure 6H), suggesting that PE contacts may pre-determine the choice of their future target genes for activation.

## DISCUSSION

In this study, we have combined high-resolution chromosomal interaction profiling with small-molecule perturbations and CRISPR targeting to investigate the dynamics, properties and potential determinants of chromosomal contacts engaging poised enhancers upon the naive-to-primed transition of hPSCs, an *in vitro* model that uniquely captures human peri-implantation epiblast development. Studying this developmental window has enabled us to temporally resolve the acquisition of poising and connectivity of human developmental enhancers. We find diverse dynamics of enhancer contacts relative to the acquisition of poising and show that PRC2 depletion early in the transition, but not inhibition of its H3K27me3 catalytic activity, weakens subsets of PE contacts. Finally, we show evidence that PE contacts persist upon developmental enhancer activation and may support gene induction.

The adaptation of low-input Capture Hi-C to enrich for PE-anchored chromatin contacts has enabled us to profile the dynamics of PE connectivity globally and at high resolution across the naive-to-primed transition. The sequence-based enrichment used in this assay has enabled profiling these contacts irrespective of their chromatin state (i.e., before and after their poising, as well as after their subsequent activation), which is challenging with techniques that enrich for contacts associated with specific recruited factors or histone modifications, such as H3K27me3 HiChIP. Consequently, our PECHi-C analysis of the naïve-to-primed transition has enabled mapping the full spectrum of PE contact dynamics in the temporally-resolved model of human peri-implantation development.

The diverse temporal patterns of PE connectivity, including contacts that are either acquired or lost before and after the onset of poising, point to multiple underlying mechanisms. It was previously shown that the contacts of several candidate PEs were lost in PRC2-knockout mouse ESCs (Cruz-Molina et al. 2017), but it remained unclear if PRC2 is involved in the establishment of new PE contacts or if it only contributes to their maintenance later in development, and how widespread its role is in either process. Our PRC2 perturbation analysis early in the naive-to-prime transition demonstrates a role for this complex in the establishment of large subsets of PE contacts in human development. Furthermore, we find that PEs engaged in the transition-invariant (‘constant’) contacts were enriched for CTCF binding in both naive and primed hPSCs, potentially implicating cohesin/CTCF-mediated loop extrusion in facilitating these contacts, consistent with findings for some PEs in mouse (Crispatzu et al. 2021). This hypothesis is further supported by the general invariance of cohesin-associated CTCF-CTCF loops between naive and primed hPSCs (Ji et al. 2016). While the large-scale chromosomal topologies mediated by Polycomb and loop extrusion, respectively, are likely in competition (Rhodes et al. 2020), it was suggested that Polycomb may also stabilise focused cohesin/CTCF-mediated contacts (Mitter et al. 2020). This hypothesis may explain why the connectivity of enhancers engaging in ‘constant’ contacts was also weakened by PRC2 depletion. Finally, we detect multitudes of transient and lost contacts during the naive-to-primed transition, which engage both developmental and intracellular transport genes. This finding parallels the recent detection of transient enhancer-promoter contacts at a later stage of development, in a mouse model of early lineage commitment (Mahara et al. 2024).

Notably, our data indicate that the effect of PRC2 on PE connectivity is likely independent of its H3K27me3 catalytic activity, since it was selectively observed upon PROTAC-mediated PRC2 degradation, but not with the catalytic inhibition of PRC2’s H3K27 methyltransferase, even though both of these perturbations have removed H3K27me3 from the majority of PEs. This finding is paralleled by PRC2’s documented role in chromatin compaction, which is retained in the absence of the methyltransferase cofactor SAM (Margueron et al. 2008; Grau et al. 2021). Catalytically-independent roles of chromatin co-factors are an emerging area of active research (Morgan and Shilatifard 2023).

The relatively mild global effects of PRC2 depletion on PE connectivity indicate that other mechanisms also play a role, which may include factors such as Polycomb Repressive Complex 1 (PRC1) activity, as well as CTCF/cohesin-mediated loop extrusion. Probing the role of PRC1 in our genetically unperturbed system is challenging due to the lack of effective small molecules targeting this complex in human. However, previous studies strongly implicate PRC1 in 3D genome organisation (Freire-Pritchett et al. 2017; Boyle et al. 2020; Crispatzu et al. 2021), and it is therefore likely that PRC1 and PRC2 act synergistically at these loci. Finally, MLL3/4 complexes that generate the H3K4me1 mark may also support PE connectivity, consistent with findings in mouse (Crispatzu et al. 2021; Kubo et al. 2024). In addition, a recent study has identified a catalytically-independent role for the H3K4me3-associated MLL2 in facilitating chromatin loops in mouse ESCs (Steindel et al. 2025). Advances in protein degradation technologies, including PROTACs and degron systems such as those established for PRC1 in mouse ESCs (Dobrinić et al. 2021), offer promise for establishing the role of these complexes upon the naive-to-primed transition in human pluripotency in the future.

By tracking the fate of PE contacts upon hPSCs differentiation, we find that many of them are retained when the respective enhancers are developmentally activated, swapping the Polycomb-associated H3K27me3 mark for H3K27 acetylation. This is consistent with findings in mouse (Crispatzu et al. 2021; Pachano et al. 2021) and recent observations that chromatin activation can reinforce looping between Polycomb-bound loci (Paldi et al. 2024). Notably, we show that these effects are largely selective for the ‘constant’ and ‘gained-early’ classes of PE contacts that are detectable in the first days of the naive-to-prime transition. This suggests that the pairing of these enhancers with their appropriate target genes is established early in human development, either before or alongside the acquisition of poising. In contrast, PE contacts that are gained at the late stages of naive-to-primed transition generally dissolve upon enhancer activation. The function of this class of contacts, including its potential involvement in embryonic lineages other than DE and NE, remains to be elucidated. The depletion of ‘gained-late’ contacts at high TF-occupancy PEs and their preferential engagement of metabolic genes not detected for other contact classes may offer initial insights for these investigations.

Finally, we find that targeting CRISPR activation machinery to a PE results in the induction of a gene connected to it through a ‘gained-early’ contact (*DLX1*), without affecting an unconnected gene located at a shorter genomic distance (*DLX2*) within the same TAD. While the CRISPRa approach is artificial, it offers the advantage of rapidly delivering targeted activating signals to PEs, modelling the very initial steps of gene induction that are largely intractable using developmental systems (Cruz-Molina et al. 2017; Crispatzu et al. 2021) and avoiding the potentially confounding effects of the global changes that occur during differentiation. Although our study only examined one locus, the fact that PE-targeted CRISPRa selectively induced the already connected gene, but not an unconnected gene in the immediate vicinity, suggests that contacts formed by enhancers in the poised state may preselect their target gene choice for future developmental activation. These effects potentially implicate a direct role for enhancer poising and Polycomb recruitment in developmental gene activation. Validating these hypotheses further requires systems to disrupt and generate PE contacts. These experiments remain challenging but are becoming more feasible with modern tools, such as CLoUD9 and LALD systems (Morgan et al. 2017; Kim et al. 2019).

In conclusion, our work presents a global, high-resolution resource of PE connectivity during human naive-to-primed transition, a critical developmental window that is unresolvable in faster-paced mouse systems. We reveal diverse dynamics of enhancer connectivity in relation to the acquisition of the poised state, and its possible molecular determinants. Our integrative analyses and perturbation experiments provide evidence for mechanisms underpinning the establishment of PE contacts prior to or during the naive-to-primed transition, and suggest a role for these contacts in facilitating gene induction upon developmental enhancer activation during lineage commitment. These findings advance our understanding of the ontogenetic origins of enhancer–promoter connectivity and target gene choice, and open new avenues to explore the role of PE dynamics in the functional transitions of pluripotency and in human enhancer-mediated pathologies.

## RESOURCE AVAILABILITY

### Lead contact

Requests for further information and resources should be directed to and will be fulfilled by the lead contact, Mikhail Spivakov.

### Material availability

This study did not generate new reagents or stable cell lines.

### Data and code availability

Sequencing data generated in this study have been uploaded to GEO. CUT&Tag data for the naive-to-primed transition are available at GEO. All the data will be made publicly available on paper acceptance. Processed datasets are listed in Tables S1-S4, with larger tables released on Open Science Framework (https://osf.io/evcy6/). Data analysis scripts will be deposited on GitHub upon paper acceptance.

## ACKNOWLEDGEMENTS

The authors would like to thank Andrew Bassett (Wellcome Sanger Institute, Cambridge) for advice on CRISPRa experimental design. We thank Laurence Game and Ivan Andrew (MRC LMS Genomics and Bioinformatics Facilities), and the members of the Babraham Institute Genomics Facility, Bioinformatics Group and Flow Cytometry Facility for technical assistance. Research in M.S. lab is core-funded by the Medical Research Council (MRC) of the UK (MC-A652-5QA20). The work was further supported by BBSRC (BB/CCG2210/1; BB/IDG2210/1; BBS/E/B/000C0522) and Wellcome (225839/Z/22/Z) to P.R-G. and MRC (MR/V02969X/1) to P.R-G. and M.R.

## AUTHOR CONTRIBUTIONS

Design and conceptualisation: M.C.N., M.D.R., M.R., P.R.-G., M.S. Formal analysis: M.C.N, M.D.R, A.A.M., S.A., M.S. Funding acquisition: P.R.-G., M.S. Investigation: M.C.N., M.D.R., A.A.M., G.L., I.S., A.B., S.E., M.R. Methodology: M.D.R., H.R.J, V.M, M.R, M.S. Project administration: P.R.-G., M.S. Resources: R.T., M.K., A.B., C.I.S., M.R., P.R.-G., M.S. Supervision: M.C.N., M.R., P.R.-G., M.S. Visualisation: M.C.N, P.R.-G., M.S. Writing – original draft: M.C.N. and M.S. Writing – review & editing: M.C.N., M.R., P.R.-G., M.S., with contributions from all authors.

## MATERIALS & METHODS

### Experimental methods

#### Cell lines

The WA09 (H9) conventional primed hPSCs (hPSCReg ID: WAe009-A) were obtained from WiCell (Thomson et al. 1998). The HNES1 naive hPSCs (hPSCReg ID: CAMe001-A; (Guo et al. 2016) were kindly shared by Prof. A. Smith. The inducible CRISPR-activation (iCRISPRa) WTC11 human iPSCs are described in (Tian et al. 2021). The use of hPSCs for these experiments is approved by the Steering Committee of the UK Stem Cell Bank (SCSC11-58 and SCSC21-01) and by the Human Research Committee at the Babraham Institute, Cambridge (HRP0021).

All cells were grown at 37 °C, 5 % CO_2_. The HNES1 naive cells were grown as described below (“Naive-to-primed hPSCs transition”). The H9 cells and the WTC11 iCRISPRa cells were grown in mTeSR-E8 basal media (StemCell Technologies, Cat. 05990), in feeder-free conditions using Geltrex (1:100 dilution in DMEM-F12, ThermoFisher Scientific, Cat. A1413302) or vitronectin XF (1:1000 dilution in D-PBS 1X) (xeno-free vitronectin, StemCell Technologies, Cat. 07180) as a coating matrix.

For calibrated CUT&Tag, Drosophila Schneider 2 (S2) cells (obtained from ThermoFisher Scientific) were cultured in a non-humidified incubator at 28 °C without additional CO_2_ in normoxic conditions. S2 cells were cultured in T75 flasks in Schneider’s *Drosophila* medium (ThermoFisher Scientific) supplemented with 10% heat-inactivated FBS (Merck). The cells grew in a semi-adherent monolayer and were passaged by gently tapping the flasks and washing gently with medium, pipetting up and down to break up clumps.

### Naive hPSC culture and naive-to-primed transition

HNES1 hPSCs were maintained in the naive state as described in (Rostovskaya 2022). In brief, cells were cultured on irradiated mouse embryonic fibroblast (MEF) feeder cells in N2B27 supplemented with 1 µM PD0325901, 10 ng/ml human LIF (made in-house), 2 µM Gö6983 (Tocris Bio-Techne, Cat. 2285), and 2 µM XAV939 (Tocris Bio-Techne, Cat. 3748). N2B27 basal medium was prepared as follows: Neurobasal and DMEM-F12 in the ratio 1:1; 0.5% N2 supplement; 1 % B27 supplement (with vitamin A); 2 mM L-glutamine; 100 mM 2-mercaptoethanol (all from ThermoFisher Scientific). Cells were passaged through single-cell dissociation using TrypLE Express (Cat. 12604021, ThermoFisher Scientific). 0.5 µL/mL Geltrex (A1413302, ThermoFisher Scientific) was added to the medium during passaging. 10 µM ROCK inhibitor (Y-27632, Cat. 688000, Millipore) was added for 24 hours after replating.

Naive-to-primed transition was performed as described in (Rostovskaya et al. 2019). For this, naive hPSCs were passaged once in feeder-free conditions, adding 1 µL/mL Geltrex to the medium during passaging, otherwise as described above. Upon reaching confluency, cells were plated at a density of 1.6 x 10^4^/cm2 in the medium for naive hPSC maintenance supplemented with 10 µM ROCK inhibitor, on plates pre-coated with 1% Geltrex. After 48 h, the medium was exchanged to N2B27 supplemented with 2 µM XAV939. The medium was refreshed daily. At confluency, cells were passaged using TrypLE Express at a 1:2 ratio in the transition medium supplemented with 10 µM ROCK inhibitor for 24 h, on Geltrex pre-coated plates. The primed state was reached by day 14 of the transition. Cells were harvested for sequencing assays on day 0 (in the naive conditions), 1, 3, 5, 7, 10, and 14 of the transition.

### Neuroectoderm differentiation

Differentiation was performed with HNES1 hPSC following the naive-to-primed transition. Neuroectoderm was induced according to (Chambers et al. 2009). N2B27 basal medium was prepared as follows: Neurobasal and DMEM-F12 in the ratio 1:1; 0.5 % N2 supplement; 1 % B27 supplement (with vitamin A); 2 mM L-glutamine; 100 mM 2-mercaptoethanol (all from ThermoFisher Scientific). hPSCs were dissociated to single cells and plated at 2.5x10^5^/cm^2^ to Geltrex-coated plates (ThermoFisher Scientific, Cat. A1413302) in N2B27 supplemented with 1 mM A8301 (Tocris Bio-Techne, Cat. 2939) and 500 nM LDN193189 (Axon Medchem, Cat. 1509). 10 mM ROCK inhibitor (Y-27632, Cat. 688000, Millipore) was added for 24 h after plating. Cells were cultured in the differentiation conditions for 10 days. To obtain a pure neuroectoderm cell population, the cells were stained for the expression of CD56 (BD, Cat. 345812) and absence of EpCAM (Biolegend, Cat. 324212), and sorted using flow cytometry.

### Definitive endoderm differentiation

Differentiation was performed with HNES1 hPSC following the naive-to-primed transition. Definitive endoderm was induced according to (Loh et al. 2014). CDM2 basal medium was prepared as follows: IMDM and F12 in the ratio 1:1, 2 mM L-Glutamine, 1 % chemically defined lipid concentrate (all from ThermoFisher Scientific), 0.1 % BSA or polyvinyl alcohol, 15 mg/mL transferrin, 450 mM monothioglycerol, 0.7 mg/mL insulin (all from Sigma). hPSCs were plated at 4x10^4^/cm^2^ to Geltrex-coated plates in their culture medium with 10 mM ROCK inhibitor. After 24 h, the medium was exchanged to CDM2 basal medium supplemented with 100 ng/mL Activin A, 3 µM CHIR99021, 10 ng/mL FGF2 (all produced in house), 100 nM PI-103 (Tocris, Bio-Techne, Cat. 2930), 3 ng/mL BMP4 (Peprotech, Cat. 120-05ET), 10 µg/mL heparin (Sigma-Aldrich, H3149) for one day; then for the next two days the following supplements were applied: 100 ng/mL Activin A, 100 nM PI-103, 20 ng/mL FGF2, 250 nM LDN193189, 10 µg/mL heparin. To obtain a pure definitive endoderm cell population, the cells were stained with antibodies against CXCR4 (BD, Cat. 555974) and CD117 (RnD Bio-Techne, Cat. IC1924A) and sorted using flow cytometry.

### Flow cytometry

Cells were dissociated using Accutase and washed using PBS with 2 % FCS. Incubation with directly conjugated antibodies diluted in PBS with 2 % FCS was carried out for 30 min at 4 °C, followed by washing and resuspending in PBS.

### Plasmid construction

Single guide RNA (sgRNA) sequences targeting the poised enhancer located in the *DLX1*-*DLX2* region (PE 280801), and a control region (Scramble) were used in this study. The sgRNA targeting the *DLX1*-*DLX2* poised enhancer was designed in-house, and the Scramble control sgRNA was designed by A. Basset. Sequences of the sense strand of sgRNAs were:

pe-DLX1 sgRNA1 5′-GGGTGTTAGGATCCTGCTAG-3′,
pe-DLX1 sgRNA2 5′-CGCAGGACCTAGTGCCCTAG-3′,
pe-DLX1 sgRNA3 5′-AGGCTGGAAGTAACGCGGCC-3′,
pe-DLX1 sgRNA4 5′-ACCCTAAACAGGTTAAAGCC-3′,
pe-DLX1 sgRNA5 5′-GGTGGTCGCTGTTCCATGGA-3′,
pe-DLX1 sgRNA6 5′-GAGCGAGTTCGGGGGACGCC-3′,
pe-DLX1 sgRNA7 5′-CGTAAGAACGACTTTTTGAG-3′,
pe-DLX1 sgRNA8 5′-GACATTGTTCTCTTGGGCGG-3′,
pe-DLX1 sgRNA9 5′-CCACCAAACCGGTATTTCTC-3′,
Control scramble sgRNA 5′-AAGATGAAAGGAAAGGCGTT-3′.

sgRNA oligonucleotides were annealed and cloned individually into the pGL3-U6-sgRNA-PGK-puromycin plasmid (Addgene, #51133). Briefly, 1.5 μg of this plasmid was digested with BsaI (NEB, Cat. R0535S) at 37 °C for 50 min, dephosphorylated with 1 μL Quick CIP (NEB, Cat. M0525S) at 37 °C for 10 min, and purified with the QIAquick PCR Purification Kit (Qiagen, Cat. 28104) following manufacturer’s protocol. Oligonucleotide annealing and phosphorylation were carried out in the same reaction, starting from 100 μM of forward and reverse sgRNAs and incubation at 37 °C for 30 min and 95 °C for 5 min with T4 DNA ligase (NEB, Cat. B0202S) and T4 PNK (NEB, Cat. M0201S). Equimolar 10 μM of annealed and phosphorylated sgRNAs oligonucleotides were ligated at 16 °C overnight into 50 ng pre-digested plasmid using T4 DNA ligase. Competent Escherichia coli (Stbl3™ strain, ThermoFisher, Cat. C737303) were transformed by heat shock at 42 °C for 30 sec, followed by incubation on ice for 2 min. After a first incubation at 37 °C in SOC medium (NEB, Cat. B9020S) with shaking at 500 rpm, cells were plated on LB media + Ampicillin (100 μg/mL) and incubated at 37 °C overnight. Successfully transformed colonies were grown in 4mL LB broth + ampicillin (100 μg/mL) at 37 °C, 180 rpm overnight, and plasmid DNA was purified using the EndoFree Plasmid Maxi Kit (Qiagen, Cat. 12362).

### PRC2 perturbation with small molecules

Naive HNES1 cells were seeded at a density of 1.6 x 10^4^/cm2 on plates coated with 1 % Geltrex and cultured in PXGL conditions. After 24 h in PXGL, either 1 µM UNC1999 (a catalytic inhibitor of the EZH1/2 subunits of PRC2), 2.5 µM UNC6852 (a PROTAC degrader of the EED subunit of PRC2), or an equivalent volume of DMSO (control) was added. After 24 h, cells were withdrawn from PXGL and cultured in N2B27 medium supplemented with XAV939 supplemented with either 1 µM UNC1999, 2.5 µM UNC6852, or DMSO. This time point was designated “day 0” of the transition experiment. Medium and inhibitors were replaced every day until day 5. Cells were harvested on day 5 in triplicate for imaging, RNA-seq, and PECHi-C experiments, and in duplicate for CUT&Tag experiments.

### Inducible CRISPR activation

WTC11 iCRISPRa cells (1.6 x 10⁶) were seeded on vitronectin XF-coated (1:1000 in D-PBS) 10-cm dishes in mTeSR-E8 basal medium. Cells were transfected with 24 µg of plasmid DNA encoding either an equimolar pool of nine sgRNAs targeting the poised enhancer *DLX1* or a scrambled sgRNA control, using TransIT-LT1 reagent (Mirus Bio, MIR2300) at a ratio of 3 µl per 1 µg DNA, in mTeSR-E8 medium supplemented with 10 µM ROCK inhibitor (Y-27632; Cambridge Bioscience, # SM02-1). Twenty-four hours post-transfection, cells were selected with 1 µg/mL puromycin in mTeSR-E8 medium for 48 h. A parallel puromycin-only control (untransfected cells) was used to confirm selection efficiency. Following a 24 h recovery period in mTeSR-E8 medium, CRISPRa was induced by addition of 20 µM trimethoprim (Cambridge Bioscience, # 738-70-5) for exactly 24 h prior to harvesting for RT-qPCR (n = 4) and 4C-seq (n = 3) experiments.

### Poised enhancer Capture Hi-C

Poised enhancer Capture Hi-C (PECHi-C) was performed in biological duplicates or triplicates as described in (Ho et al. 2021; Malysheva et al. 2022; Ray-Jones et al. 2025), with minor modifications. Cells were dissociated into single-cell suspensions using TrypLE. 1 million cells per condition were fixed in 1X D-PBS (Sigma-Aldrich, Cat. D8537-500ML) containing 2 % paraformaldehyde (Electron Microscopy Sciences, Cat. 15714-S) for 10 min at room temperature, and crosslinking was quenched with incubation in 0.125 M glycine for 5 min at room temperature followed by 5 min on ice. Cells were pelleted by centrifugation, washed with PBS, and lysed for 30 min on ice in Hi-C lysis buffer (10 mM Tris-HCl pH 8.0, 10 mM NaCl, 0.2% IGEPAL CA-630, protease inhibitors). All centrifuging steps during the lysis, prior to *DpnII* digestion, were carried out in a swinging-bucket centrifuge at 4°C, 300 x *g* for 5 min.The released nuclei were permeabilised and digested overnight with DpnII (NEB, Cat. R0543L). Restriction overhangs were filled in with biotin-dATP (Jena Bioscience, Cat. NU-835-BIO14-S), and proximity ligation was performed for 4 h at 16 °C using T4 DNA ligase (Invitrogen, Cat. 15224-025). Crosslinks were reversed overnight at 65 °C with proteinase K (Roche, Cat. 3115836001), and DNA was purified with AMPure XP beads (Beckman Coulter, Cat. A63881). DNA was fragmented by tagmentation to generate ≤1,000 bp fragments and biotinylated junctions were enriched using MyOne C1 streptavidin beads (Life Technologies, Cat. 65001). Libraries were amplified directly on beads with 5 PCR cycles, purified with AMPure XP beads, and subjected to poised enhancer capture using a custom-designed Agilent SureSelect system (described below - see “Computational analysis”), following the manufacturer’s protocol. Libraries were sequenced as 150 bp paired-end reads.

### CUT&Tag

Cleavage Under Targets and Tagmentation (CUT&Tag) was performed as described by (Kaya-Okur et al. 2019) with minor modifications. The CUT&Tag data across the naïve-to-primed transition were generated in a companion publication (M.D.R., M.R. et al., in preparation). Cells were harvested on day 0 (in the naive conditions), 1, 3, 5, 7, 10, and 14 of the transition in biological triplicates. Cells were washed twice in PBS, dissociated into single-cell suspensions with TrypLE (5 min, 37 °C), neutralized, washed, and resuspended in Wash Buffer (20 mM HEPES pH 7.5, 150 mM NaCl, 0.5 mM spermidine, protease inhibitors). Concanavalin A beads (10 µL per 100,000 cells; Stratech, Cat. BP531-BAN-3ml) were activated in Binding Buffer (20 mM HEPES pH 7.5, 10 mM KCl, 10 mM CaCl₂, 10 mM MnCl₂) and incubated with 100,000 cells per reaction at room temperature for 10 min with agitation. Bead-cell complexes were resuspended in Antibody Buffer (2 mM EDTA pH 8, 0.1 % BSA in Dig-Wash Buffer) containing primary antibodies (1:100) and incubated overnight at 4 °C on a tube rotator. Included antibodies: H3K4me1 (Abcam, Cat. ab8895), H3K4me3 (Active Motif, Cat. 39159), H3K27me3 (Cell Signaling Technology, Cat. 9733), H3K27ac (Millipore, Cat. MABE647), H2AK119ub (Cell Signaling Technology, Cat. D27C4), and IgG control (Abcam, Cat. ab2410). After a wash in Dig-Wash Buffer (0.05 % digitonin in Wash Buffer), complexes were incubated with a secondary antibody (guinea pig α-rabbit IgG, 1:100; Rockland, Cat. ABIN101961) for 30 min at room temperature on a tube rotator. After three washes in Dig-Wash Buffer, bead-cell complexes were incubated with pAG-Tn5 (1:250; EpiCypher, Cat. 15-1117) in Dig300-Wash Buffer (20 mM HEPES pH 7.5, 300 mM NaCl, 0.5 mM spermidine, protease inhibitors, 0.01 % digitonin) for 1 h at room temperature. After three washes in Dig300-Wash Buffer, tagmentation was activated in Tagmentation Buffer (10 mM MgCl₂ in Dig300-Wash Buffer) for 1 h at 37 °C. The tagmentation reaction was stopped by the addition of 10 µL 0.5 M EDTA, 3 µL 10 % SDS, and 2.5 µL 20 mg/mL proteinase K, and incubated for 1 h at 55 °C. DNA was purified with 1 X AMPure XP beads (Beckman Coulter) to enrich for ≥ 100 bp fragments. Libraries were amplified using KAPA HiFi HotStart polymerase (Roche Sequencing Store, Cat. KK2601) with 11–18 PCR cycles (depending on the primary antibody used), purified with AMPure XP beads, and sequenced as 150 bp paired-end reads.

Calibrated CUT&Tag was performed as described above with the following modifications. Cells were harvested by incubation with TrypLE for 5-7 minutes at 37 °C. TrypLE was quenched by wash buffer (0.1 % BSA in DMEM/F12) and cells were collected by centrifugation at 300 x *g* for 3 min. Cells were washed in 1X PBS without Calcium or Magnesium (PBS -/-) and collected by centrifugation at 300 x *g* for 3 minutes. Cells were resuspended in PBS -/- before addition of 2X nuclear extraction buffer (25 mM HEPES pH 7.5 (Gibco, Cat. 15630), 12.5 mM Potassium Chloride, 0.25 mM Spermidine, 0.13 % Triton X-100 (Sigma, Cat. T9284), 25% Glycerol and 1 x Protease inhibitor cocktail (Roche) in water) to 1X and incubation on ice for 10 min with intermittent vortexing. Nuclei were collected by centrifugation at 1,300 x *g* for 4 min at 4 °C. Nuclei were resuspended in cold PBS -/- and fresh 16 % PFA was added to a final concentration of 0.1 % followed by rocking gently at room temperature for 2 min. Fixation was stopped by addition of 1.25 M glycine. Nuclei were collected by centrifugation at 1,300 x *g* for 4 min at 4°C and resuspended in wash buffer. Human cells (n = 20,000) were mixed with *Drosophila* cells (n = 8,000) per reaction prior to addition of concanavalin-beads. The following antibodies were used at a 1:50 dilution H3K27me3 (CST, Cat. 9733), H2AK119ub (Cell Signalling, Cat. D27C4), Rabbit anti-mouse IgG (H+L) (Invitrogen, Cat. 31188), and 1:100 Mouse (G3A) IgG (CST, Cat. 5415S). For amplification of tagmented DNA, 14 PCR cycles were used for H3K27me3, and 18 cycles were used for H2AK119ub and IgG samples.

### ChIP-seq

ChIP-seq was performed as described (Bendall and Semprich 2022) with minor modifications. Twenty million naive or primed hPSCs were harvested, cross-linked in suspension with 2 mM di(N-succinimidyl) glutarate for 45 min and 1 % methanol-free paraformaldehyde for 12.5 min, then quenched with 0.125 M glycine. Nuclei were prepared by sequential extraction and wash buffers and lysed in 25 mM Tris pH 7.5, 150 mM NaCl, 5 mM EDTA, 0.1 % Triton X-100, 1 % SDS, and 0.5 % sodium deoxycholate. The sonicated chromatin was diluted 1:10 in ChIP buffer (150 mM NaCl, 25 mM Tris pH 7.5, 5 mM EDTA, 1 % Triton X-100, 0.1 % SDS and 0.5 % Sodium Deoxycholate); 5 % was saved as input. The remainder was incubated overnight at 4 °C with 10 µg anti-KAT3B/P300 antibody (Abcam, Cat. ab14984). Protein G Dynabeads were blocked with BSA/yeast tRNA and added to capture immune complexes. Beads were washed sequentially with low-, high-, and LiCl-salt buffers, followed by TE buffer. Chromatin and inputs were eluted in 1 % SDS/0.1 M NaHCO₃ with proteinase K and RNase A, then cross-links were reversed at 65 °C. DNA was purified with AMPure XP beads (Beckman Coulter). Libraries were prepared using the NEBNext Ultra II DNA kit (NEB) with indexed adapters and sequenced as 75-bp single-end reads.

### RNA-seq on PRC2 perturbed cells

Total RNA was extracted from PRC2-perturbed cells treated with DMSO, PROTAC, or UNC1999 at day 5 of the transition, with three biological replicates per condition. Cells were washed with PBS and harvested in RLT buffer from the RNeasy Mini Kit (Qiagen, Cat. 74104) and stored at -80C. Once thawed, the suspension was passed through a QIAshredder (Qiagen, Cat. 79656) and the RNA purified using the RNeasy Mini Kit following manufacturer’s instructions. Contaminating DNA was removed using the Turbo DNA free kit (ThermoFisher Scientific, Cat. AM1907) following manufacturer’s instructions. RNA-seq libraries were prepared using the Watchmaker RNA Library Prep Kit with ribosomal RNA depletion following manufacturer’s instructions. Libraries were sequenced as 75bp paired-end reads.

### Retro-transcription and quantitative PCR (RT-qPCR)

Cells were washed twice in D-PBS, dissociated using TrypLE, and lysed in RLT buffer (from RNeasy Mini Kit, QIAGEN, Cat. 74104). Lysates were homogenized with QIAshredder columns (QIAGEN, Cat. 79654), and total RNA was extracted using the RNeasy Mini Kit according to the manufacturer’s instructions. 2 to 4 µg of RNA was reverse-transcribed with SuperScript™ IV Reverse Transcriptase (Thermo Fisher Scientific, cat. 18090200). Quantitative PCR was performed in triplicate on 200 µg cDNA in 20 µL reaction using PowerUp™ SYBR™ Green Master Mix (ThermoFisher, Cat. A25742). Primer sequences were as follows:

*DLX1* (F: 5’-TCAGTTCGGTGCAGTCCTA-3’, R: 5’-ACCGTGCTCTTCTCCGAGT-3’), *DLX2* (F: 5’-ACACCTCCTACGCTCCCTATG-3’, R: 5’-TCACTATCCGAATTTCAGGCTCA-3’), *METAP1D* (F: 5’- TTTGGCATCATGCAAACGACA-3’, R: 3’-CTGCGCCGACCTTTGATTG-5’), and *GAPDH* (F: 5’-GAAGACTGTGGATGGCCCC-3’, R: 5’-TCCACCACTGACACGTTGG-3’).

Ct values were normalised to Scrmbl-transfected cells and *GAPDH* controls. Relative expression was calculated by the 2^-ΔΔCt^ method (Livak and Schmittgen 2001). Statistical comparisons between independent biological replicates (n = 4 per condition) were performed using a Wilcoxon rank-sum test with continuity correction, applying the normal approximation due to tied values.

### 4C-seq

The viewpoint region (location: chr2:172,096,016-172,096,312) within the poised enhancer (location: chr2:172,095,731-172,096,369) in the *DLX1-DLX2* region was chosen according to the (Krijger et al. 2020) guidelines, as well as primary and secondary restriction enzymes and reading and non-reading primers.

Reading primer: location: chr2:172,096,288-172,096,308; sequence: 5’-TACACGACGCTCTTCCGATCTTCCCAAATTCGCTCCATCAGT-3’.

Non-reading primer: location: chr2:172,096,038-172,096,057; sequence: 5’-ACTGGAGTTCAGACGTGTGCTCTTCCGATCTCCCCGAACTCGCTCAAAAGT-3’.

All 4C-seq experiments were performed in three independent biological replicates. Approximately 10 million cells were washed in PBS, dissociated with TrypLE, and crosslinked in 2 % formaldehyde (Fisher bioreagents, Cat. BP531-500) for 10 min, as described by (Krijger et al. 2020). Crosslinking was quenched with 0.13 M glycine, and nuclei were isolated in lysis buffer (50 mM Tris-HCl, pH 7.5, 150 mM NaCl, 0.5 % IGEPAL CA-630, 1 % TritonX-100, 5 mM EDTA, and protease inhibitors) for 20 min. Chromatin was digested overnight with 200 U *DpnII* (NEB, Cat. R0543T), the primary restriction enzyme, then overnight ligated with 50 U of DNA ligase (Invitrogen Cat. 15224017) following published protocols (Krijger et al. 2020; Taylor et al. 2022; Ray-Jones et al. 2025). Crosslink were reversed with proteinase K, and 3C templates were purified with AMPure XP beads (Beckman Coulter, Cat. A63881).

Up to 60 μg of template was diluted to 100 ng/μL, and digested overnight with *NlaIII* (NEB, #R0125S, 1 U/μg DNA template), the secondary restriction enzyme, followed by a second overnight ligation at a final DNA concentration of 5 ng/μL using 2 U/μg T4 DNA ligase (Invitrogen). DNA was purified by Phenol Chloroform Isoamyl Alcohol (P:C:I) extraction and AMPure XP beads, according to (Ray-Jones et al. 2025).

4C libraries were generated in two rounds of PCR, as described in (Krijger et al. 2020). Briefly, first-round PCRs (16 cycles) were performed with Expand Long Template polymerase (Roche, Cat. 11681842001) on four 200-ng reactions per sample, followed by pooling and 0.8 X AMPure XP beads cleanup to remove long primers. A second PCR (12 cycles) was performed with up to 100 ng of template per reaction to introduce Illumina sequencing adapters. Finally, libraries were purified as above and sequenced as 60-bp paired-end reads.

### Western blot

Protein samples were denatured in 1x SDS loading buffer at 95 °C for 5 min. Proteins (10-30 μg) were separated on SDS-PAGE gels then transferred onto nitrocellulose membranes. Blocking, incubations and washes used 5% milk powder in PBS with 0.1% Tween-20. Primary antibodies were H3K27me3 (Cell Signaling Technology; 1:1000), H4 (Cell Signaling Technology; 1:1000), EED (Cell Signaling Technology; 1:1000), and EZH2 (Cell Signaling Technology; 1:1000).

### Computational analyses

#### Design of the Poised Enhancer Capture Hi-C system

PEs were identified based on their chromatin profile and the joint presence of H3K4me1 and H3K27me3 signals. For this purpose, six published ChIP-seq datasets were used (Bernstein et al. 2010; Rada-Iglesias et al. 2011; Vallot et al. 2015; Luo et al. 2020) in combination with CUT&RUN data for H3K4me1 (GEO IDs GSM4798822, GSM4798823, GSM4798822 from (Chovanec et al. 2021)) and H3K27me3 (released with this study) in H9 primed hESCs. Raw sequencing files were aligned to the GRCh38 reference genome using bowtie2 (v2.3.5) (Langmead and Salzberg 2012). Peaks were called using Macs2 peak-caller (v2.1.1), (Zhang et al. 2008)), with the broad peak option used for H3K27me3. ChIPQC Bioconductor R package (v.1.21.0, (Carroll et al. 2014)) was used to compute quality metrics of the ChIPseq datasets and those with a FRiP (Fraction of Reads in Peak) 10% were excluded from the analyses. P-value cutoffs were determined based on the precision-recall analysis versus H9 histone modification data presented in (Rada-Iglesias et al. 2011), either directly or integrated using ChromHMM (Ernst and Kellis 2012) in (Freire-Pritchett et al. 2017), resulting in the p-value cutoffs of 1e-6.5 for ChIP-seq and p-value = 1e-5 for CUT&RUN datasets, respectively. The final capture design included bivalent regions identified by both ChIP-seq and CUT&RUN, except for those overlapping annotated gene promoters, which were excluded. The identified PE regions were mapped to the *DpnII*-digested genome fragments. Biotinylated RNA probes complementary to these regions were designed for each fragment, resulting in a total of 32,142 *DpnII*-digested genome fragments included in the capture system design. RNA probes were then generated and provided by Agilent Technologies (SureSelect CustomDNA Target Enrichment Probes, 6.0 - 11.999 Mbp). The GRCh38 coordinates of all and PE-harbouring (baited) *DpnII* fragments are released on OpenScienceFramework (https://osf.io/evcy6/).

### Generation of the metaplots for naive and primed cells based on public datasets

Enhancer sets were filtered for coverage in the range of 100-450 reads per 20kb windows for all plotted datasets. Aligned ATAC-seq (GSM2698915, GSM2698916, GSM2698917, GSM2698918, GSM2698919, GSM2698920, GSM2698921, GSM2698922 from (Pastor et al. 2018)), CUT&RUN, ChIP-seq and Input (GSM2698940, GSM2698941, GSM2698942, GSM2901723, GSM2901724, GSM2901725 from (Pastor et al. 2018)) BAM files for H3K27me3 (GSM5569915, GSM5569916, GSM5569923, GSM5569924 from (Zijlmans et al. 2022)), H3K4me1 (GSM4798820; GSM4798821, GSM4798822, GSM4798823, GSM4798824 from (Chovanec et al. 2021)), and p300 ChIP-seq (this study), were converted to bigwig files using bamCoverage from Deeptools v3.5.4 using default parameters. BigWig files for replicates of the same condition were combined using bigwigAverage with default parameters. The bigwig files for each enhancer set and modality, plus input samples, were used to generate a matrix file using computeMatrix reference-point based on the centre of each enhancer +/-10kb, adding the option --missingDataAsZero. Plots were created from matrices using plotHeatmap with manually defined scales for both the colourmap and heatmap to ensure consistency between plots for the same modality but different enhancer subsets.

### Primary processing and visualisation of PECHi-C data

Read processing, alignment, and filtering were performed using the Hi-C User Pipeline (HiCUP, (Wingett et al. 2015), modified to consider both across- and within-read ligation junctions (https://github.com/StevenWingett/HiCUP/tree/combinations, v0.7.4, (Freire-Pritchett et al. 2021)). CHiCAGO input files were generated using chicagoTools v1.30.0 (Cairns et al. 2016) for bins of *DpnII* fragments spanning ∼5 kb, leaving the captured (‘baited’) PE-harbouring fragments unbinned to increase resolution. CHiCAGO (v1.30.0) was run as previously described (Cairns et al. 2016; Freire-Pritchett et al. 2021). The ‘other ends’ of detected contacts were annotated with transcription start sites downloaded from GENCODE (v.36 (Frankish et al. 2021)). CHiCAGO results were summarised across samples using *makePeakMatrix.R* from ChicagoTools with default parameters. The full peak matrices are released on OpenScienceFramework (https://osf.io/evcy6/), and those filtered for PE-promoter and PE-TSS contacts provided in Tables S1 (naive-to-primed transition) and S3A (PRC2 perturbations).

To visualise the contacts of the candidate PEs (baitIDs 1340 and 580854) in naive and primed hPSCs, *plotBaits*() function from the Chicago R package was used with a maximum interaction distance of 500 kb. Enrichment of PE-interacting regions for histone modifications was estimated using the *peakEnrichment4Features*() function in CHiCAGO. Locus-specific visualisations combining PECHi-C and CUT&Tag signals were generated using the *plotgardener* R package (v1.6.4; (Kramer et al. 2022)).

### Primary processing and visualisation of CUT&Tag data

Aligned BAM files were sorted, indexed and biological triplicates were merged with *samtools* (v1.20, (Danecek et al. 2021)). For visualization, both merged and individual BAM files were converted to bigWig format using *bamCoverage* from *deepTools* (v3.5.6, (Ramírez et al. 2016)), and signal tracks were visualized with *plotgardener* (v1.6.4). For integration with PECHi-C data, we generated feature count matrices for each dataset on the bins of *DpnII* fragments used in PECHi-C analysis (with the PE-harbouring ‘baited’ fragments unbinned) and cross-normalised them using size factors produced by *DESeq2’s* estimateSizeFactors (v1.40.2, (Love et al. 2014). The pileup matrices are available on OpenScienceFramework (https://osf.io/evcy6/). For analyses in Figures 3AB, RPKM values at the baited fragments were computed by normalising read counts to fragment length.

### Sparse correlation analysis of PECHi-C signals

To assess correlations between PECHi-C datasets across the HNES1 naive-to-primed transition and H9 primed hPSCs, CHiCAGO interaction scores from each cell type/time point were used as input for the Sparse Correlations for Compositional data analysis (SparCC; (Friedman and Alm 2012)) with the *SpiecEasi* R package (v0.1.5). The resulting correlation matrices were visualised as heatmaps, with a colour gradient representing correlation strength.

### Annotation of poised enhancers with enhancer activity across tissues

PE coordinates were overlapped with the annotations of active candidate cis-regulatory elements (cCREs) from the ENCODE consortium’s ‘enhancer decoration matrix’ (https://downloads.wenglab.org/cCRE_decoration.matrix.1.gz; (Rozowsky et al. 2023)). From this matrix, cCREs were filtered to retain 635,906 cCREs described as “active” in at least one tissue. The genomic coordinates of cCREs were downloaded from UCSC Genome Browser (https://genome.ucsc.edu/cgi-bin/hgTrackUi?db=hg38&g=encodeCcreCombined). Sample-level annotations were combined at tissue level, and the results were visualised as histograms using *ggplot2* in R.

### Poised enhancer annotation in VISTA enhancer database

Experimentally validated enhancers were retrieved from the VISTA Enhancer Browser (Visel et al. 2007; Kosicki et al. 2025) and filtered to retain only positively validated elements (curation_status = “positive”) with human-specific identifiers (prefix “hs”). Genomic overlaps between VISTA enhancers and poised enhancer baits were computed in R using the *data.table* package *foverlaps* function (Barrett et al. 2006) (v1.17.6). Counts of overlapping enhancers per tissue were summarised and visualised as horizontal bar plots using the R package *ggplot2* (v3.5.2).

### Temporal clustering of PE contacts

To investigate dynamic changes in chromatin contacts during the HNES1 naive-to-primed transition experiment, we applied clustering using the Dynamic Time Warping dissimilarity metric implemented in the tsclust function of the *dtwclust* R package v5.5.12 (Alexis Sarda-Espinosa 2015). The analysis focused on 9,998 contacts between PEs and ‘other ends’ containing either annotated TSS or other PEs that had CHiCAGO scores above 5 in any generated dataset (including the naive-to-primed transition time course, H9 primed hPSCs, and NE- and DE-differentiated samples) and above 3 in at least one time point in the transition. Partitionings into 3 to 8 clusters were tested with the *tsclust* function, and the final clustering into 6 clusters was chosen based on inspection of cluster trajectories. Clusters 2 and 6 were merged post-hoc, resulting in five biologically interpretable temporal classes of contacts: ‘constant’ (cluster 3), ‘gained early’ (cluster 1), ‘gained late’ (clusters 2 and 6), ‘transient’ (cluster 5), and ‘lost’ (cluster 4). The final clusters are listed in Table S1. Combinations of temporal classes of contact per PE were visualised using an UpSet plot (*UpSetR* package v1.4.0 (Lex et al. 2014)).

### Clustering of chromatin features at poised enhancers involved in interaction

H3K27me3 spreading was estimated by counting the number of 10bp bins with more than 25 read counts in the 20-kb vicinity of PE centres in the H3K27me3 CUT&Tag data in primed cells. The binding profiles of TFs and co-factors were based on the following public and in-house datasets in naive and primed hPSC: DPPA2 and DPPA4 (ChIP-seq; A. Malcolm, P.R.G et al., in revision); CTCF (ChIP-seq; (Ji et al. 2016)); OCT4 ((Ji et al. 2016)); NANOG ((Chovanec et al. 2021; Osnato et al. 2021)); SOX2 ((Chovanec et al. 2021; Osnato et al. 2021)); TFAP2C ((Pastor et al. 2018; Dong et al. 2020)); Mediator ((Ji et al. 2016)); BRM ((Huang et al. 2021)); BRG1 ((Huang et al. 2021)); BAF155 ((Huang et al. 2021)). For each of these datasets, we obtained read pileups at the PE-harbouring baited *DpnII* fragments and normalised them for library size using DESeq2’s *estimateScalingFactorsForMatrix()*. Values were transformed using the inverse hyperbolic sine (arcsinh), scaled and submitted to K-means clustering (k = 5). The resulting clusters (listed in Table S2) were annotated according to their characteristic enrichment patterns: ‘Broad H3K27me3’, ‘CTCF + TFs’, ‘Intermediate’, ‘High Occupancy’ and ‘Low Occupancy’. Heatmaps of row-normalised signal intensities were generated using the R package *ppclust* (v1.1.0.1, (Cebeci 2019)).

### Enrichment of PE clusters for temporal classes of contacts

To assess the association between temporal chromatin dynamics (‘constant,’ ‘gained early,’ ‘gained late,’ ‘transient,’ and ‘lost’) and poised enhancer baits clustering (‘Broad H3K27me3’, ‘CTCF + TFs’, ‘Intermediate’, ‘High Occupancy’ and ‘Low Occupancy’), we used a Chi-square test on the 5x5 contingency table (*chisq.test*() in R). We visualised this contingency table as a heatmap of log-odds ratios:

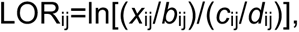

where *x*_ij_ is the count in cell *(i,j)*, *b*_ij_=∑_m!=j_*x*_im_, *c*_ij_=∑_k!=i_*x*_kj_, *d*_ij_=∑_k!=i,m!=j_*x*_km_. The heatmap was generated using the *ggplot2* package (v3.5.2) in R. To highlight significant enrichment and depletion compared with random expectation, we denoted absolute chi-square standardised residuals above 1.96 (returned by *chisq.test()*) with asterisks on the heatmap.

### Functional annotation of contacted genes detected by PECHi-C

Gene Ontology (GO) enrichment analysis for each temporal class was performed using *clusterProfiler* (v4.8.2, (Wu et al. 2021)). Enrichment for Biological Process terms was assessed with Benjamini-Hochberg multiple testing correction. To compare enrichment patterns across temporal classes, enrichment results were concatenated, and the top 20 GO terms per condition (ranked by adjusted *p*-value) were visualised using *clusterProfiler’s dotplot*(). GO terms were further grouped by semantic similarity using *GOSemSim* (v2.26.1, (Yu 2020)) with the Wang method, followed by hierarchical clustering. GO results were visualised in a multi-condition dot plot, colour-coded by four semantic clusters.

### Network analysis of PE contact rewiring upon naive-to-primed transition

PE baits were filtered to include the PE baits that contain at least one ‘gained early’ or ‘gained late’, as well as one ‘lost’ contact. The R package *igraph* (v2.0.3) was used to construct an adjacency matrix from all contacts of these PEs (PE-TSS and PE-PE) and annotated them by contact class. The full network was visualised using *igraph.plot*() with the Kamada-Kawai layout algorithm. For the ‘zoomed in’ subnetwork, we subsampled 50 random disjoint subgraphs and supplemented them with all detected contacts with *HOX* genes, removing duplicated edges. The ‘zoomed in’ subnetwork was visualised using the R package *ggraph* with the Fruchterman-Reingold force-directed layout algorithm, with the density of *HOX* gene nodes per subnetwork colour-coded as yellow halos.

### PRC2 perturbation experiments

#### RNA-seq data

Read quality control and mapping to the GRCh38 genome were performed by the Babraham Institute bioinformatics platform according to their pipeline. BAM files were sorted and indexed with *samtools* (v1.20), and gene-level quantification was performed using *featureCounts* (*Subread* package, v2.0.8, (Liao et al. 2014)) with default parameters and Gencode v47 annotation (Frankish et al. 2023). Only unique mapped reads overlapping exonic regions were counted. Output count matrices from all samples (DMSO, PROTAC- and UNC1999-treated) were merged into a single combined matrix. Differential expression analysis was performed using *DESeq2* (v1.40.2), with a design that accounted for both treatment and biological replicate (∼ condition + replicate). DESeq2 normalised the counts, estimated gene-wise dispersion, and fitted a negative binomial generalised linear model. Comparisons of interest included PROTAC-perturbated versus DMSO and UNC1999-perturbated versus DMSO. Genes with an adjusted p-value (Benjamini-Hochberg correction) < 0.05 were considered statistically significant. Upregulated and downregulated genes were categorised based on log2 fold change, with |log2FC| ≥ 0.1 indicating small effect changes. Significantly differentially expressed genes were annotated with HGNC gene symbols using the org.Hs.eg.db annotation database (v3.17.0 (Carlson 2017), AnnotationDbi v1.62.2 (Hervé Pagès 2017)). The expression analysis table is released on OpenScienceFramework: (https://osf.io/evcy6/). Volcano plots were generated with *ggplot2*, colour-coding genes as upregulated, downregulated, or not significant, and labelling only highly significant genes (padj < 1e-3 and |log2FC| > 0.41).

#### CUT&Tag data

Primary processing of calibrated CUT&Tag samples was carried out as follows. FastQ files were trimmed with TrimGalore (v0.6.5,https://www.bioinformatics.babraham.ac.uk/projects/trim_galore/), followed by mapping to either GRCh38 for human reads or BDGP6 for spike-in reads using a Nextflow implementation of bowtie2 with the additional settings –very-sensitive -I 10 - X 700 for both genomes. Aligned BAM files for each genome were then quality filtered (MapQ > 20), sorted and indexed using samtools (v1.21). Scale factors were calculated as follows : 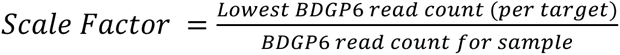.

GRCh38-aligned BAM files were converted to reference-calibrated bigWig files using *deepTools bamCoverage* (v3.5.6, (Ramírez et al. 2016)) scaling by these scale factors.

For quantitative analysis, individual calibrated H3K27me3 bigWig files were converted into pileUp files with features corresponding to baited PE fragments using *bwtool’s summary* function (v20170428, (Pohl and Beato 2014)). The pileup files were released on OpenScienceFramework (https://osf.io/evcy6/). The resulting datasets were tested for differential counts (each perturbation versus DMSO) using the R package *DESeq2*. Regions with adjusted p-value < 0.05 and negative log2 fold change were considered significantly reduced in H3K27me3 occupancy upon perturbation, whereas regions with adjusted *p*-value > 0.5 were defined as non-responsive. To identify genomic regions commonly affected by both PROTAC and UNC1999 perturbations (shown in Table S3B), we intersected the sets of significantly depleted H3K27me3 regions from each comparison. The full analysis table is available on OpenScienceFramework (https://osf.io/evcy6/). For the density profiles and heatmap visualisations, individual replicate bigWig files were merged using *deepTools bigwigAverage*. These merged files for H3K27me3 and H2AK119ub were used across the three experimental conditions to generate signal matrices with the *computeMatrix* function from *deepTools* in the reference-point mode with the centre of poised enhancers (±10 kb) as reference and the --skipZeros option. Heatmaps and average profile plots of the signal distributions were produced using the *plotHeatmap* and *plotProfile* functions of *deepTools*, respectively.

#### PECHi-C data

Raw PECHi-C data for the three conditions (three replicates each of DMSO, PROTAC, and UNC1999) were aligned using *HiCUP combinations* and chicagoTools as described above (“Primary processing and visualisation of PECHi-C data”). Significant contacts (listed in Table S3A) were detected by CHiCAGO on datasets pooled across replicates for each condition and visualised using R package *plotGardener*. For differential analysis (summarised in Table S3B), two replicates each for PROTAC and UNC1999 were merged to balance the sequencing depths across replicates. The read counts table used for this analysis is available on OpenScienceFramework (https://osf.io/evcy6/). Total connectivity by PE bait in each replicate was computed using a custom R script, and changes in connectivity upon each PRC2 perturbation were estimated using *DESeq2*.

#### Integrative analysis of PECHi-C and CUT&Tag

Distributions of DESeq2 test statistics for each perturbation were visualised using R package *ggplot2* (v3.5.1), separately for PEs exhibiting a significant loss of H3K27me3 (‘perturbed PEs’) versus not (‘intact PEs’) in the respective perturbation. The differences between DESeq2 test statistic distributions (*t-*values) for H3K27me3-perturbed and intact PE pools were assessed using the Wilcoxon rank-sum test. The same analysis was repeated separately for the subsets of PEs engaging contacts from four temporal classes: ‘constant’, ‘gained early’, ‘transient’, and ‘lost’; the ‘gained late’ class was excluded as these contacts were generally absent at day 5 (annotated in Table S3B). Single-sample Wilcoxon tests against the *t-*value of zero were used to assess the overall reduction in connectivity of perturbed PEs, and pairwise Wilcoxon rank-sum tests with Bonferroni correction (*rstatix* v0.7.2 (Kassambara 2019)) were used to evaluate differences between perturbed and intact PEs. Finally, to assess whether changes in chromosomal contacts upon PROTAC treatment depend on temporal contact dynamics, we restricted the analysis to 1,590 PE baits engaging only one of the four classes of contacts and assessed class-specific differences in PE connectivity changes (expressed as DESeq2 test statistics) using the Kruskal-Wallis test. Boxplots for all analyses were produced using the R package *ggplot2*.

### Analysis of PE chromatin state and contact dynamics after NE and DE differentiation

For each PE fragment, we obtained CUT&Tag read counts for H3K27me3, H3K4me1, H3K4me3 and H3K27ac. We then used standard DESeq2 analysis on this data matrix to detect differential signals for each histone modification between day 14 of the transition and DE, and between day 14 and NE (DESeq2 adjusted p-value<0.05). Based on these data for H3K27me3, H3K4me1 and H3K27ac, we assigned chromatin state dynamics to each PE upon NE and DE differentiation (Table S4). PEs undergoing developmental activation, which were the focus of this analysis, were defined as “H3K27me3 down, H3K27ac up”. For PEs showing these dynamics, we generated the boxplots of the distribution of log2-fold changes in arcsinh-Chicago scores upon NE and DE differentiation for each temporal class of contacts. The “Lost”, “Transient” and *de novo* contacts in differentiated cells were combined into the “Other” category. Log2-fold changes were computed using a pseudocount of 0.1.

### 4C-seq data processing and visualisation

Sequencing reads were trimmed with *fastp* v0.23.4 (Chen et al. 2018) requiring a minimal length of 45 bp. Only read 1 from paired-end sequencing was used for downstream analysis. Processed reads were mapped and filtered with *pipe4C* (Krijger et al. 2020) (https://github.com/deLaatLab/pipe4C) using the following parameters: minimum mapping quality of 1, retention of uniquely mapped reads, and cis-only analysis. The genomic coordinate chr2:172,096,280 was used as the viewpoint position. Quality control of individual experiments was based on the diagnostic reports generated by *pipe4C*. Interaction profiles were exported as WIG files and visualised in the *Integrative Genomics Viewer* (IGV) (Robinson et al. 2011). TAD boundaries in the *DLX1*-*DLX2* region were based on public HiC data on WTC11 cells (dataset 4DNFI2YA1USU downloaded from the 4DN data portal https://data.4dnucleome.org/ (Bertero et al. 2019)).

---

**Figure S1:**
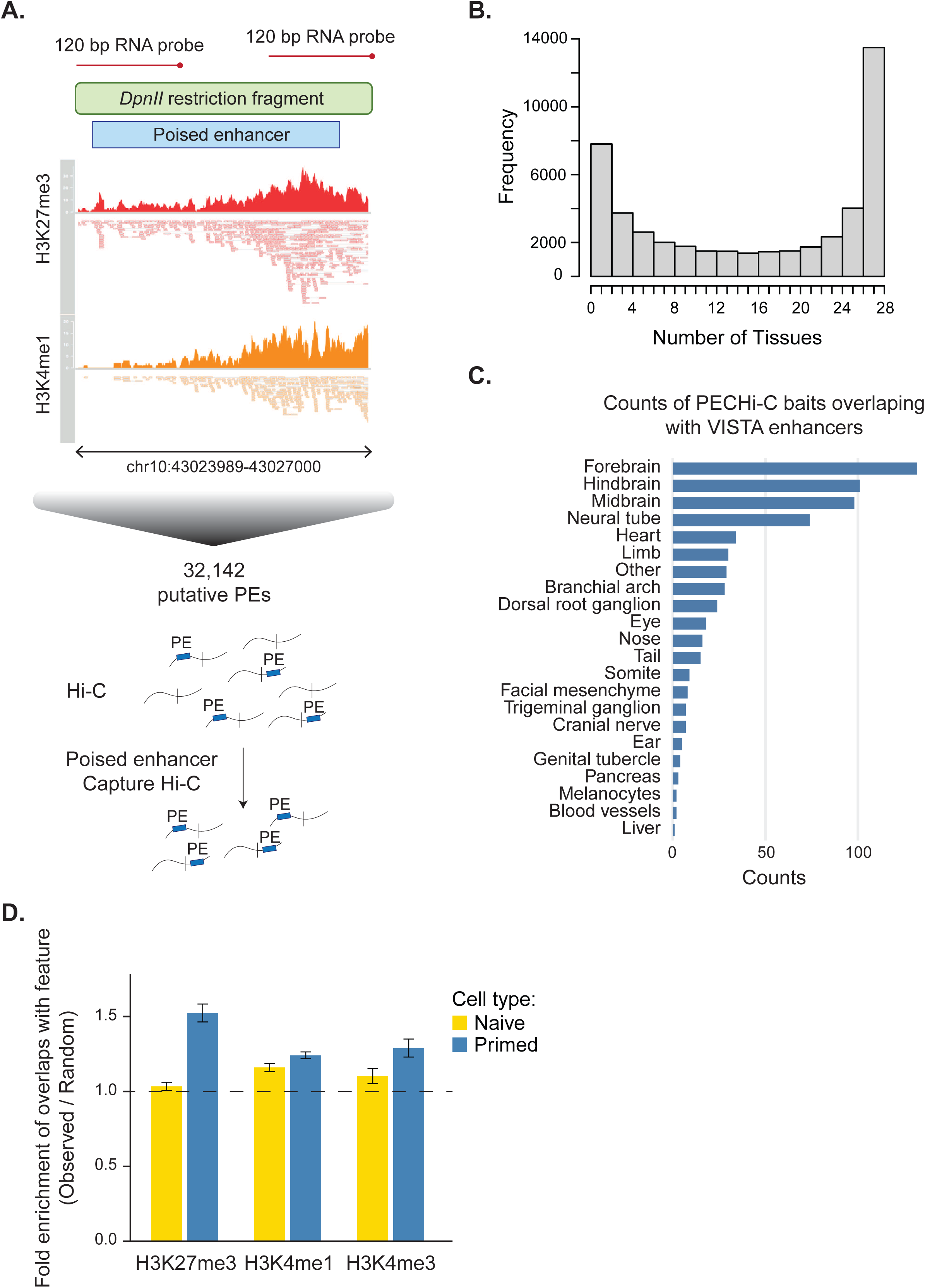
Additional information on PE Capture Hi-C analysis in naive and primed hPSCs. (*A*) Schematic of the Poised Enhancer Capture Hi-C (PECHi-C) experiment. Public H3K27me3 (red) and H3K4me1 (orange) ChIP-seq datasets from primed hPSCs were used to identify 32,142 PEs. The flanks of the *DpnII* fragments harbouring these PEs were used to design a sequence capture library of 120bp biotinylated RNA baits to enrich Hi-C material for contacts involving (at least on one end) these baited fragments. (*B*) Histogram showing the number of hPSC PEs found in the “active” state in human adult tissues according to the ENCODE project. (*C*) Bar plot showing the number of PEs with *in vivo* tissue-specific activity overlapping poised enhancers according to VISTA Enhancer Browser, grouped by annotated tissue activity. (*D*) Enrichment of PE-contacted regions for histone marks versus distance-matched random regions in naive (yellow bars) and primed cells (blue bars). Error bars represent 95% confidence intervals over 100 random samples. The enrichment is statistically significant where the baseline of one (dashed line) lies below the entirety of the 95% confidence intervals.

**Figure S2:**
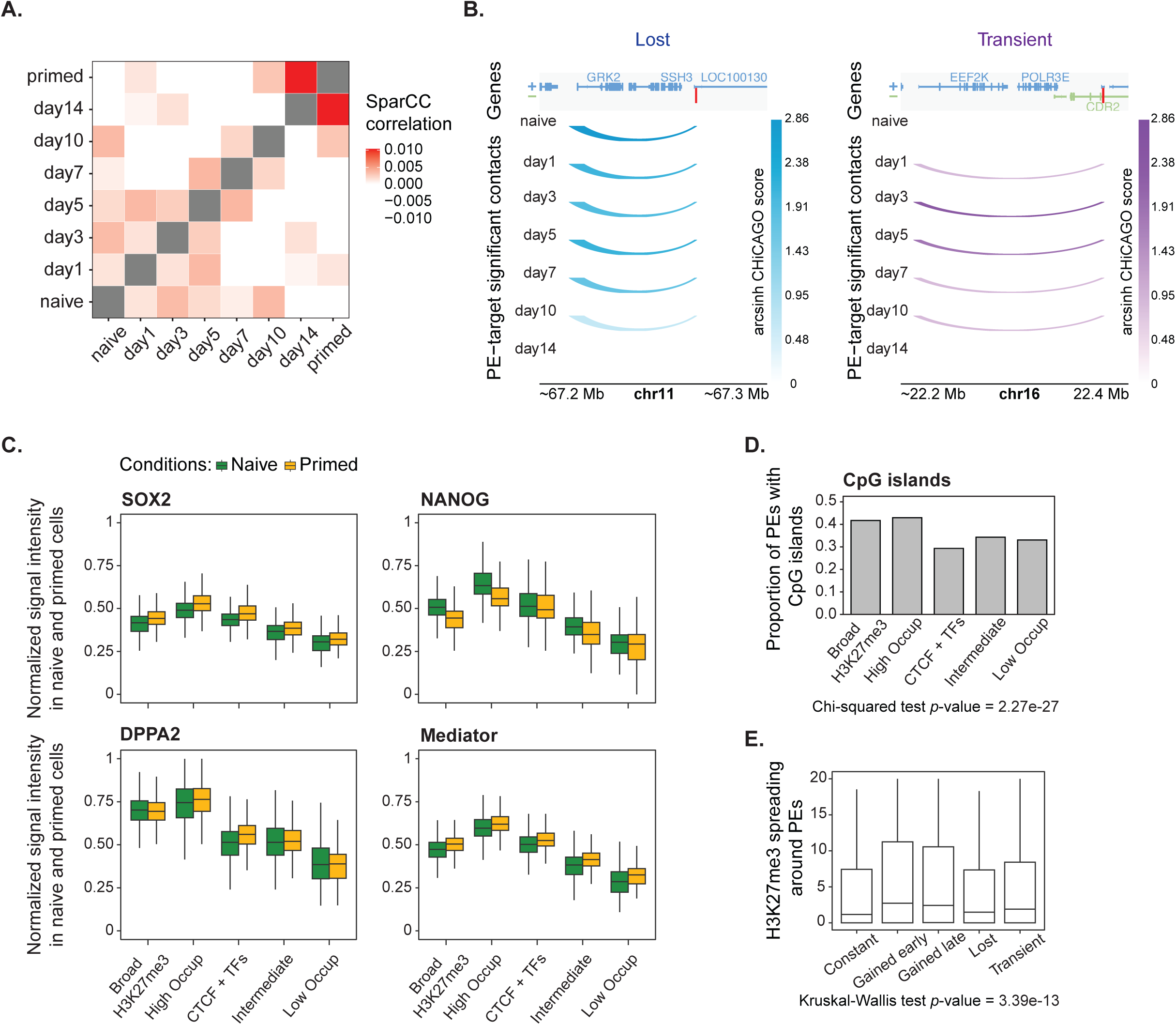
Additional information on PE contact dynamics during the naive-to-primed hPSC transition. (*A*) Heatmap showing SparCC values (sparse correlations for compositional data) for pairwise relationships between PECHi-C contact scores in transitioning and primed cells (the grey colour indicates a correlation of 1). (*B*) Representative examples of chromatin contacts detected by PECHi-C: a contact between PE 88624 (red bar) and the *GRK2* promoter during the naive-to-primed transition mapped to the ‘lost’ temporal class, and a ‘transient’ contact between PE 188917 and the *EEF2K* promoter. The arcs’ principal colour reflects temporal class annotation from Figure 2B and their colour intensities represent arcsinh-transformed CHiCAGO scores (see colour key). Absence of an arc corresponds to the CHiCAGO score of zero. (+) and (-) denote gene strandedness. (*C*) Normalised ChIP signal intensities for SOX2, NANOG, DPPA2, and Mediator at PE clusters in naive (green) and primed (yellow) cells. Note that as these data were used in cluster definition, conventional statistical testing for their differences between clusters is not appropriate due to “double-dipping”. (*D*) Proportion of PEs containing CpG islands in the different classes of poised enhancers. (*E*) The spreading of H3K27me3 around PEs involved in different temporal contact dynamics.

**Figure S3:**
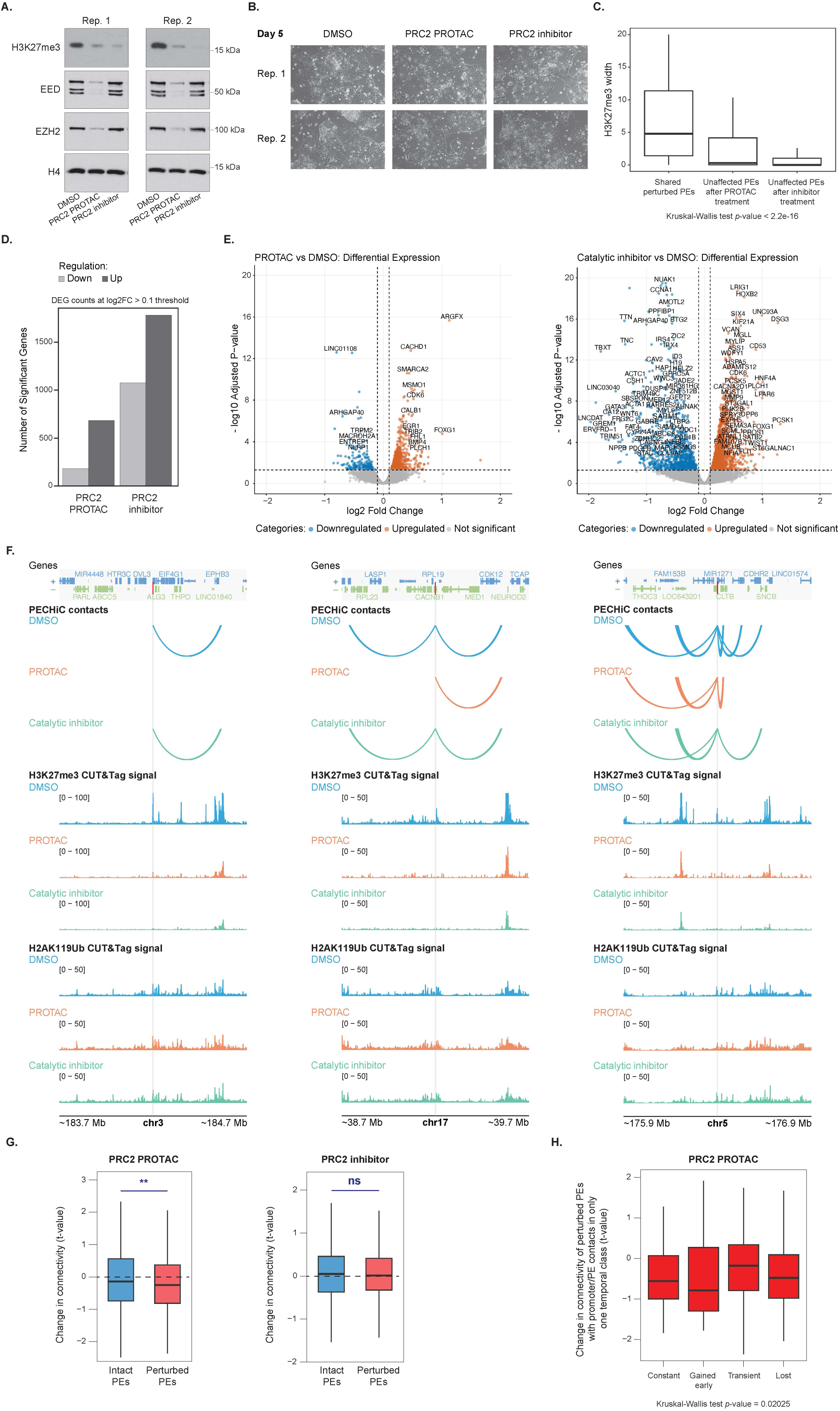
Additional information on PRC2 perturbation analysis. (A)Western blot analysis of protein extracts from hPSCs at day 5 of transition treated with a PRC2 PROTAC degrader, a PRC2 catalytic inhibitor, or DMSO control for 5 days. The blots were probed for H3K27me3, EED, EZH2, and H4 antibodies (n = 2).(B) Images of hPSCs at day 5 of transition treated with the same compounds (n = 2).(C) Barplots showing the spreading of H3K27me3 around PEs perturbed by both PROTAC and catalytic inhibitor treatment (“shared”), or unperturbed following either PROTAC or catalytic inhibitor treatment. (*D*) Number of significantly differentially expressed genes (|log₂FC| ≥ 0.1, adjusted p-value < 0.05) measured by RNA-seq after PRC2 PROTAC or PRC2 catalytic inhibitor treatment relative to DMSO control. Light grey bars: significantly downregulated genes; dark grey bars: significantly upregulated genes. (*E*) Volcano plots of differential gene expression after PRC2 PROTAC (left) or PRC2 catalytic inhibitor (right) treatment, compared to DMSO. Blue dots indicate significantly downregulated genes (padj < 0.05 & log₂FC ≤ -0.1); orange dots indicate significantly upregulated genes (padj < 0.05 & log₂FC ≥ 0.1); the rest of the genes are shown as grey dots. Dashed grey lines indicate significance and fold-change thresholds. Labels are shown for genes with padj < 1e−3 and |log₂FC| > 0.41, corresponding to a ∼30% change in expression. (*F*) Additional examples of loss of contacts detected by PECHi-C (top), alongside the H3K27me3 (middle) and H2AK119Ub (bottom) CUT&Tag signals in the respective loci following treatment with a PRC2 PROTAC (orange) or a PRC2 catalytic inhibitor (green), compared to DMSO (blue). Grey and red bars highlight signals at PE baits. (*G*) Changes in the overall connectivity of intact (blue) and perturbed (red) PEs after treatment with PRC2 PROTAC or PRC2 catalytic inhibitor. (*H*) Change in enhancer connectivity for perturbed PE baits engaged in promoter/PE contact(s) of only a single temporal class. (*G, H*): Y axis shows PE-level *t*-values for the difference in normalised read counts between treatment and DMSO control samples estimated by DESeq2.

**Figure S4:**
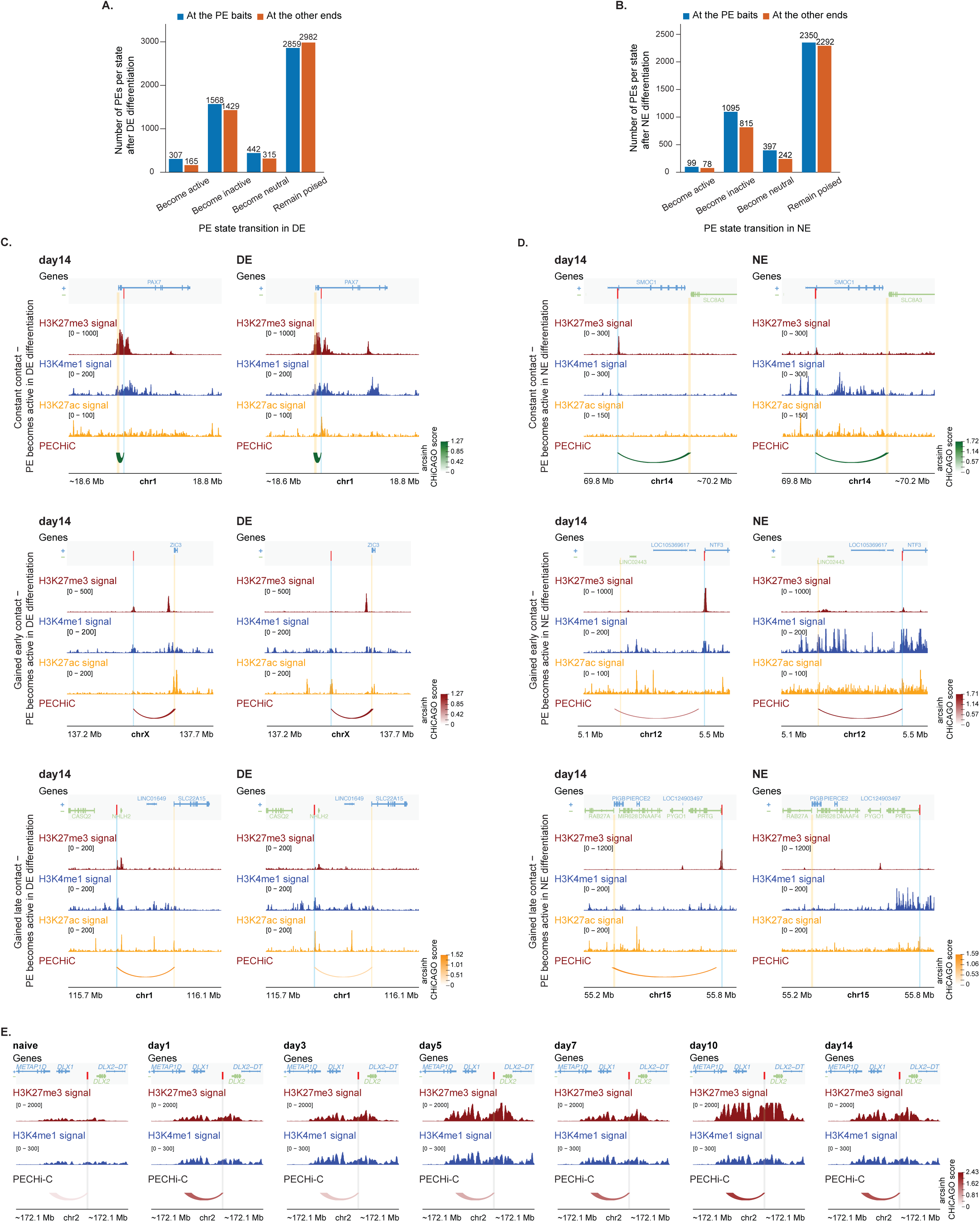
Additional information on the dynamics of PE contacts after developmental or ectopic activation. (*A,B*) Number of poised enhancers (PEs) (blue bars) and contacted other ends (promoters or other PEs; orange bars) for different chromatin state dynamics upon differentiation of hPSCs into definitive endoderm (DE) (*A*) or neuroectoderm (NE) (*B*) following the naive-to-primed transition : ‘become active’ (significant loss of H3K27me3 and gain of H3K27ac); ‘become inactive’ (significant loss of H3K4me1 and no gain of H3K27ac); ‘become neutral’ (significant loss of H3K27me3 and H3K4me1, no gain of H3K27ac); ‘remain poised’ (no change in all three histone marks). (*C, D*) Representative examples of PE contacts showing different temporal dynamics upon the naive-to-primed transition (top: ‘constant’, middle: ‘gained early’, bottom: ‘gained late’) on day 14 of the transition and after differentiation into DE (*C*) or NE (*D*). H3K27me3 (dark red), H3K4me1 (dark blue), and H3K27ac (orange) CUT&Tag signals shown in the panels below, followed by arcs representing PECHi-C contacts between a PE and a gene promoter or another PE within the indicated genomic region. The arcs’ principal colour reflects temporal class annotation from Figure 2B and their colour intensities represent arcsinh-transformed CHiCAGO scores (see colour key). Absence of an arc corresponds to the CHiCAGO score of zero. Red and light blue bars highlight signals at PE baits, and yellow bars highlight signals at their contacted promoters/PEs. (*E*) Visualisation of the H3K27me3 (dark red) and H3K4me1 (dark blue) CUT&Tag signals and the ‘gained-early’ PECHi-C contact (red arcs) between PE 280801 (red and grey bars) and the *DLX1* promoter during the naive-to-primed transition, with arc color intensity reflecting arcsinh-transformed CHiCAGO scores.

## SUPPLEMENTAL TABLES

**Table S1.** Significant PE-promoter and PE-PE contacts detected during the naive-to-primed transition of naive hPSCs, DE and NE differentiation, as well as in primed hPSCs, and PE temporal classes defined on their basis.

**Table S2.** Enrichment of chromatin features at PEs and PE chromatin clusters defined on their basis.

**Table S3.** Supporting data for PRC2 perturbation analysis. (A) Significant PE-promoter and PE-PE contacts detected in untreated and PRC2 PROTAC- or inhibitor-treated hPSCs at day 5 of naive-to-primed transition. (B) Changes in PE connectivity upon either treatment.

**Table S4.** Dynamics of PE contacts and histone modifications upon DE and NE differentiation.

## REFERENCES

Agostinho de Sousa J, Wong C-W, Dunkel I, Owens T, Voigt P, Hodgson A, Baker D, Schulz EG, Reik W, Smith A, et al. 2023. Epigenetic dynamics during capacitation of naïve human pluripotent stem cells. Sci Adv 9: eadg1936.

Alexis Sarda-Espinosa. 2015. dtwclust: Time Series Clustering Along with Optimizations for the Dynamic Time Warping Distance. 6.0.0. https://CRAN.R-project.org/package=dtwclust.

Azuara V, Perry P, Sauer S, Spivakov M, Jørgensen HF, John RM, Gouti M, Casanova M, Warnes G, Merkenschlager M, et al. 2006. Chromatin signatures of pluripotent cell lines. Nat Cell Biol 8: 532–538.

Barrett T, Dowle M, Srinivasan A, Gorecki J, Chirico M, Hocking T, Schwendinger B, Krylov I. 2006. data.table: Extension of ‘data.framè. 1.17.8. https://CRAN.R-project.org/package=data.table.

Bendall A, Semprich CI. 2022. Chromatin Profiling of Human Naïve Pluripotent Stem Cells. In Human Naïve Pluripotent Stem Cells (ed. P. Rugg-Gunn), Vol. 2416 of Methods in Molecular Biology, pp. 181–200, Springer US, New York, NY https://link.springer.com/10.1007/978-1-0716-1908-7_12.

Bernstein BE, Mikkelsen TS, Xie X, Kamal M, Huebert DJ, Cuff J, Fry B, Meissner A, Wernig M, Plath K, et al. 2006. A Bivalent Chromatin Structure Marks Key Developmental Genes in Embryonic Stem Cells. Cell 125: 315–326.

Bernstein BE, Stamatoyannopoulos JA, Costello JF, Ren B, Milosavljevic A, Meissner A, Kellis M, Marra MA, Beaudet AL, Ecker JR, et al. 2010. The NIH Roadmap Epigenomics Mapping Consortium. Nat Biotechnol 28: 1045–1048.

Bertero A, Fields PA, Ramani V, Bonora G, Yardimci GG, Reinecke H, Pabon L, Noble WS, Shendure J, Murry CE. 2019. Dynamics of genome reorganization during human cardiogenesis reveal an RBM20-dependent splicing factory. Nat Commun 10: 1538.

Boroviak T, Loos R, Lombard P, Okahara J, Behr R, Sasaki E, Nichols J, Smith A, Bertone P. 2015. Lineage-Specific Profiling Delineates the Emergence and Progression of Naive Pluripotency in Mammalian Embryogenesis. Dev Cell 35: 366–382.

Boyle S, Flyamer IM, Williamson I, Sengupta D, Bickmore WA, Illingworth RS. 2020. A central role for canonical PRC1 in shaping the 3D nuclear landscape. Genes Dev 34: 931–949.

Cairns J, Freire-Pritchett P, Wingett SW, Várnai C, Dimond A, Plagnol V, Zerbino D, Schoenfelder S, Javierre B-M, Osborne C, et al. 2016. CHiCAGO: robust detection of DNA looping interactions in Capture Hi-C data. Genome Biol 17: 127.

Calo E, Wysocka J. 2013. Modification of enhancer chromatin: what, how, and why? Mol Cell 49: 825–837.

Carlson M. 2017. org.Hs.eg.db. https://bioconductor.org/packages/org.Hs.eg.db.

Carroll TS, Liang Z, Salama R, Stark R, de Santiago I. 2014. Impact of artifact removal on ChIP quality metrics in ChIP-seq and ChIP-exo data. Front Genet 5: 75.

Cebeci Z. 2019. Comparison of Internal Validity Indices for Fuzzy Clustering. J Agric Inform 10. http://journal.magisz.org/index.php/jai/article/view/537.

Chambers SM, Fasano CA, Papapetrou EP, Tomishima M, Sadelain M, Studer L. 2009. Highly efficient neural conversion of human ES and iPS cells by dual inhibition of SMAD signaling. Nat Biotechnol 27: 275–280.

Chen S, Zhou Y, Chen Y, Gu J. 2018. fastp: an ultra-fast all-in-one FASTQ preprocessor. Bioinforma Oxf Engl 34: i884–i890.

Chovanec P, Collier AJ, Krueger C, Várnai C, Semprich CI, Schoenfelder S, Corcoran AE, Rugg-Gunn PJ. 2021. Widespread reorganisation of pluripotent factor binding and gene regulatory interactions between human pluripotent states. Nat Commun 12: 2098.

Coussement L, Ciarchi M, Kolmogorova A, Meyer TD, Rulands S, Reik W, Rostovskaya M. 2025. A transcriptional clock of the human pluripotency transition. http://biorxiv.org/lookup/doi/10.1101/2025.03.13.643129.

Creyghton MP, Cheng AW, Welstead GG, Kooistra T, Carey BW, Steine EJ, Hanna J, Lodato MA, Frampton GM, Sharp PA, et al. 2010. Histone H3K27ac separates active from poised enhancers and predicts developmental state. Proc Natl Acad Sci 107: 21931–21936.

Crispatzu G, Rehimi R, Pachano T, Bleckwehl T, Cruz-Molina S, Xiao C, Mahabir E, Bazzi H, Rada-Iglesias A. 2021. The chromatin, topological and regulatory properties of pluripotency-associated poised enhancers are conserved in vivo. Nat Commun 12: 4344.

Cruz-Molina S, Respuela P, Tebartz C, Kolovos P, Nikolic M, Fueyo R, van Ijcken WFJ, Grosveld F, Frommolt P, Bazzi H, et al. 2017. PRC2 Facilitates the Regulatory Topology Required for Poised Enhancer Function during Pluripotent Stem Cell Differentiation. Cell Stem Cell 20: 689–705.e9.

Dahl JA, Jung I, Aanes H, Greggains GD, Manaf A, Lerdrup M, Li G, Kuan S, Li B, Lee AY, et al. 2016. Broad histone H3K4me3 domains in mouse oocytes modulate maternal-to-zygotic transition. Nature 537: 548–552.

Danecek P, Bonfield JK, Liddle J, Marshall J, Ohan V, Pollard MO, Whitwham A, Keane T, McCarthy SA, Davies RM, et al. 2021. Twelve years of SAMtools and BCFtools. GigaScience 10: giab008.

Dimitrova E, Feldmann A, van der Weide RH, Flach KD, Lastuvkova A, de Wit E, Klose RJ. 2022. Distinct roles for CKM-Mediator in controlling Polycomb-dependent chromosomal interactions and priming genes for induction. Nat Struct Mol Biol 29: 1000–1010.

Dobrinić P, Szczurek AT, Klose RJ. 2021. PRC1 drives Polycomb-mediated gene repression by controlling transcription initiation and burst frequency. Nat Struct Mol Biol 28: 811–824.

Dong C, Beltcheva M, Gontarz P, Zhang B, Popli P, Fischer LA, Khan SA, Park K, Yoon E-J, Xing X, et al. 2020. Derivation of trophoblast stem cells from naïve human pluripotent stem cells. eLife 9: e52504.

Du Z, Zheng H, Kawamura YK, Zhang K, Gassler J, Powell S, Xu Q, Lin Z, Xu K, Zhou Q, et al. 2020. Polycomb Group Proteins Regulate Chromatin Architecture in Mouse Oocytes and Early Embryos. Mol Cell 77: 825–839.e7.

Eckersley-Maslin MA, Parry A, Blotenburg M, Krueger C, Ito Y, Franklin VNR, Narita M, D’Santos CS, Reik W. 2020. Epigenetic priming by Dppa2 and 4 in pluripotency facilitates multi-lineage commitment. Nat Struct Mol Biol 27: 696–705.

Ernst J, Kellis M. 2012. ChromHMM: automating chromatin-state discovery and characterization. Nat Methods 9: 215–216.

Frankish A, Carbonell-Sala S, Diekhans M, Jungreis I, Loveland JE, Mudge JM, Sisu C, Wright JC, Arnan C, Barnes I, et al. 2023. GENCODE: reference annotation for the human and mouse genomes in 2023. Nucleic Acids Res 51: D942–D949.

Frankish A, Diekhans M, Jungreis I, Lagarde J, Loveland JE, Mudge JM, Sisu C, Wright JC, Armstrong J, Barnes I, et al. 2021. GENCODE 2021. Nucleic Acids Res 49: D916–D923.

Freire-Pritchett P, Ray-Jones H, Della Rosa M, Eijsbouts CQ, Orchard WR, Wingett SW, Wallace C, Cairns J, Spivakov M, Malysheva V. 2021. Detecting chromosomal interactions in Capture Hi-C data with CHiCAGO and companion tools. Nat Protoc 16: 4144–4176.

Freire-Pritchett P, Schoenfelder S, Várnai C, Wingett SW, Cairns J, Collier AJ, García-Vílchez R, Furlan-Magaril M, Osborne CS, Fraser P, et al. 2017. Global reorganisation of cis-regulatory units upon lineage commitment of human embryonic stem cells. eLife 6: e21926.

Friedman J, Alm EJ. 2012. Inferring correlation networks from genomic survey data. PLoS Comput Biol 8: e1002687.

Grau D, Zhang Y, Lee C-H, Valencia-Sánchez M, Zhang J, Wang M, Holder M, Svetlov V, Tan D, Nudler E, et al. 2021. Structures of monomeric and dimeric PRC2:EZH1 reveal flexible modules involved in chromatin compaction. Nat Commun 12: 714.

Gretarsson KH, Hackett JA. 2020. Dppa2 and Dppa4 counteract de novo methylation to establish a permissive epigenome for development. Nat Struct Mol Biol 27: 706–716.

Guo G, von Meyenn F, Santos F, Chen Y, Reik W, Bertone P, Smith A, Nichols J. 2016. Naive Pluripotent Stem Cells Derived Directly from Isolated Cells of the Human Inner Cell Mass. Stem Cell Rep 6: 437–446.

Gupta RM, Hadaya J, Trehan A, Zekavat SM, Roselli C, Klarin D, Emdin CA, Hilvering CRE, Bianchi V, Mueller C, et al. 2017. A Genetic Variant Associated with Five Vascular Diseases Is a Distal Regulator of Endothelin-1 Gene Expression. Cell 170: 522–533.e15.

Heintzman ND, Hon GC, Hawkins RD, Kheradpour P, Stark A, Harp LF, Ye Z, Lee LK, Stuart RK, Ching CW, et al. 2009. Histone modifications at human enhancers reflect global cell-type-specific gene expression. Nature 459: 108–112.

Heinz S, Romanoski CE, Benner C, Glass CK. 2015. The selection and function of cell type-specific enhancers. Nat Rev Mol Cell Biol 16: 144–154.

Hervé Pagès MC. 2017. AnnotationDbi. https://bioconductor.org/packages/AnnotationDbi.

Ho JSY, Mok BW-Y, Campisi L, Jordan T, Yildiz S, Parameswaran S, Wayman JA, Gaudreault NN, Meekins DA, Indran SV, et al. 2021. TOP1 inhibition therapy protects against SARS-CoV-2-induced lethal inflammation. Cell 184: 2618–2632.e17.

Huang J, Liu X, Li D, Shao Z, Cao H, Zhang Y, Trompouki E, Bowman TV, Zon LI, Yuan G-C, et al. 2016. Dynamic Control of Enhancer Repertoires Drives Lineage and Stage-Specific Transcription during Hematopoiesis. Dev Cell 36: 9–23.

Huang X, Park K, Gontarz P, Zhang B, Pan J, McKenzie Z, Fischer LA, Dong C, Dietmann S, Xing X, et al. 2021. OCT4 cooperates with distinct ATP-dependent chromatin remodelers in naïve and primed pluripotent states in human. Nat Commun 12: 5123.

Javierre BM, Burren OS, Wilder SP, Kreuzhuber R, Hill SM, Sewitz S, Cairns J, Wingett SW, Várnai C, Thiecke MJ, et al. 2016. Lineage-Specific Genome Architecture Links Enhancers and Non-coding Disease Variants to Target Gene Promoters. Cell 167: 1369–1384.e19.

Ji X, Dadon DB, Powell BE, Fan ZP, Borges-Rivera D, Shachar S, Weintraub AS, Hnisz D, Pegoraro G, Lee TI, et al. 2016. 3D Chromosome Regulatory Landscape of Human Pluripotent Cells. Cell Stem Cell 18: 262–275.

Kassambara A. 2019. rstatix: Pipe-Friendly Framework for Basic Statistical Tests. 0.7.2. https://CRAN.R-project.org/package=rstatix.

Kaya-Okur HS, Wu SJ, Codomo CA, Pledger ES, Bryson TD, Henikoff JG, Ahmad K, Henikoff S. 2019. CUT&Tag for efficient epigenomic profiling of small samples and single cells. Nat Commun 10: 1930.

Kempfer R, Pombo A. 2020. Methods for mapping 3D chromosome architecture. Nat Rev Genet 21: 207–226.

Kim JH, Rege M, Valeri J, Dunagin MC, Metzger A, Titus KR, Gilgenast TG, Gong W, Beagan JA, Raj A, et al. 2019. LADL: light-activated dynamic looping for endogenous gene expression control. Nat Methods 16: 633–639.

Kojima Y, Tam OH, Tam PPL. 2014. Timing of developmental events in the early mouse embryo. Semin Cell Dev Biol 34: 65–75.

Kosicki M, Baltoumas FA, Kelman G, Boverhof J, Ong Y, Cook LE, Dickel DE, Pavlopoulos GA, Pennacchio LA, Visel A. 2025. VISTA Enhancer browser: an updated database of tissue-specific developmental enhancers. Nucleic Acids Res 53: D324–D330.

Kramer NE, Davis ES, Wenger CD, Deoudes EM, Parker SM, Love MI, Phanstiel DH. 2022. Plotgardener: cultivating precise multi-panel figures in R. Bioinforma Oxf Engl 38: 2042–2045.

Krijger PHL, Geeven G, Bianchi V, Hilvering CRE, de Laat W. 2020. 4C-seq from beginning to end: A detailed protocol for sample preparation and data analysis. Methods San Diego Calif 170: 17–32.

Kubo N, Chen PB, Hu R, Ye Z, Sasaki H, Ren B. 2024. H3K4me1 facilitates promoter-enhancer interactions and gene activation during embryonic stem cell differentiation. Mol Cell 84: 1742–1752.e5.

Langmead B, Salzberg SL. 2012. Fast gapped-read alignment with Bowtie 2. Nat Methods 9: 357–359.

Lex A, Gehlenborg N, Strobelt H, Vuillemot R, Pfister H. 2014. UpSet: Visualization of Intersecting Sets. IEEE Trans Vis Comput Graph 20: 1983–1992.

Liao Y, Smyth GK, Shi W. 2014. featureCounts: an efficient general purpose program for assigning sequence reads to genomic features. Bioinforma Oxf Engl 30: 923–930.

Liu X, Wang C, Liu W, Li J, Li C, Kou X, Chen J, Zhao Y, Gao H, Wang H, et al. 2016. Distinct features of H3K4me3 and H3K27me3 chromatin domains in pre-implantation embryos. Nature 537: 558–562.

Livak KJ, Schmittgen TD. 2001. Analysis of relative gene expression data using real-time quantitative PCR and the 2(-Delta Delta C(T)) Method. Methods San Diego Calif 25: 402–408.

Loh KM, Ang LT, Zhang J, Kumar V, Ang J, Auyeong JQ, Lee KL, Choo SH, Lim CYY, Nichane M, et al. 2014. Efficient endoderm induction from human pluripotent stem cells by logically directing signals controlling lineage bifurcations. Cell Stem Cell 14: 237–252.

Love MI, Huber W, Anders S. 2014. Moderated estimation of fold change and dispersion for RNA-seq data with DESeq2. Genome Biol 15: 550.

Luo Y, Hitz BC, Gabdank I, Hilton JA, Kagda MS, Lam B, Myers Z, Sud P, Jou J, Lin K, et al. 2020. New developments on the Encyclopedia of DNA Elements (ENCODE) data portal. Nucleic Acids Res 48: D882–D889.

Lupiáñez DG, Kraft K, Heinrich V, Krawitz P, Brancati F, Klopocki E, Horn D, Kayserili H, Opitz JM, Laxova R, et al. 2015. Disruptions of Topological Chromatin Domains Cause Pathogenic Rewiring of Gene-Enhancer Interactions. Cell 161: 1012–1025.

Mahara S, Prüssing S, Smialkovska V, Krall S, Holliman S, Blum B, Dachtler V, Borgers H, Sollier E, Plass C, et al. 2024. Transient promoter interactions modulate developmental gene activation. Mol Cell 84: 4486–4502.e7.

Malysheva V, Ray-Jones H, Cazares TA, Clay O, Ohayon D, Artemov P, Wayman JA, Rosa MD, Petitjean C, Booth C, et al. 2022. High-resolution promoter interaction analysis in Type 3 Innate Lymphoid Cells implicates Batten Disease gene CLN3 in Crohn’s Disease aetiology. bioRxiv 2022.10.19.512842.

Margueron R, Li G, Sarma K, Blais A, Zavadil J, Woodcock CL, Dynlacht BD, Reinberg D. 2008. Ezh1 and Ezh2 Maintain Repressive Chromatin through Different Mechanisms. Mol Cell 32: 503–518.

Mifsud B, Tavares-Cadete F, Young AN, Sugar R, Schoenfelder S, Ferreira L, Wingett SW, Andrews S, Grey W, Ewels PA, et al. 2015. Mapping long-range promoter contacts in human cells with high-resolution capture Hi-C. Nat Genet 47: 598–606.

Mitter M, Gasser C, Takacs Z, Langer CCH, Tang W, Jessberger G, Beales CT, Neuner E, Ameres SL, Peters J-M, et al. 2020. Conformation of sister chromatids in the replicated human genome. Nature 586: 139–144.

Morgan MAJ, Shilatifard A. 2023. Epigenetic moonlighting: Catalytic-independent functions of histone modifiers in regulating transcription. Sci Adv 9: eadg6593.

Morgan SL, Mariano NC, Bermudez A, Arruda NL, Wu F, Luo Y, Shankar G, Jia L, Chen H, Hu J-F, et al. 2017. Manipulation of nuclear architecture through CRISPR-mediated chromosomal looping. Nat Commun 8: 15993.

Nakamura T, Okamoto I, Sasaki K, Yabuta Y, Iwatani C, Tsuchiya H, Seita Y, Nakamura S, Yamamoto T, Saitou M. 2016. A developmental coordinate of pluripotency among mice, monkeys and humans. Nature 537: 57–62.

Noordermeer D, Leleu M, Schorderet P, Joye E, Chabaud F, Duboule D. 2014. Temporal dynamics and developmental memory of 3D chromatin architecture at Hox gene loci. eLife 3: e02557.

Osnato A, Brown S, Krueger C, Andrews S, Collier AJ, Nakanoh S, Quiroga Londoño M, Wesley BT, Muraro D, Brumm AS, et al. 2021. TGFβ signalling is required to maintain pluripotency of human naïve pluripotent stem cells. eLife 10: e67259.

Pachano T, Sánchez-Gaya V, Ealo T, Mariner-Faulí M, Bleckwehl T, Asenjo HG, Respuela P, Cruz-Molina S, Muñoz-San Martín M, Haro E, et al. 2021. Orphan CpG islands amplify poised enhancer regulatory activity and determine target gene responsiveness. Nat Genet 53: 1036–1049.

Paldi F, Szalay MF, Di Stefano M, Jost D, Reboul H, Cavalli G. 2024. Transient histone deacetylase inhibition induces cellular memory of gene expression and three-dimensional genome folding. http://biorxiv.org/lookup/doi/10.1101/2024.11.21.624660.

Pastor WA, Liu W, Chen D, Ho J, Kim R, Hunt TJ, Lukianchikov A, Liu X, Polo JM, Jacobsen SE, et al. 2018. TFAP2C regulates transcription in human naive pluripotency by opening enhancers. Nat Cell Biol 20: 553–564.

Pohl A, Beato M. 2014. bwtool: a tool for bigWig files. Bioinforma Oxf Engl 30: 1618–1619.

Potjewyd F, Turner A-MW, Beri J, Rectenwald JM, Norris-Drouin JL, Cholensky SH, Margolis DM, Pearce KH, Herring LE, James LI. 2020. Degradation of Polycomb Repressive Complex 2 with an EED-Targeted Bivalent Chemical Degrader. Cell Chem Biol 27: 47–56.e15.

Rada-Iglesias A, Bajpai R, Swigut T, Brugmann SA, Flynn RA, Wysocka J. 2011. A unique chromatin signature uncovers early developmental enhancers in humans. Nature 470: 279–283.

Ramírez F, Ryan DP, Grüning B, Bhardwaj V, Kilpert F, Richter AS, Heyne S, Dündar F, Manke T. 2016. deepTools2: a next generation web server for deep-sequencing data analysis. Nucleic Acids Res 44: W160–W165.

Ray-Jones H, Spivakov M. 2021. Transcriptional enhancers and their communication with gene promoters. Cell Mol Life Sci 78: 6453–6485.

Ray-Jones H, Sung CK, Chan LT, Haglund A, Artemov P, Della Rosa M, Ruje L, Burden F, Kreuzhuber R, Litovskikh A, et al. 2025. Genetic coupling of enhancer activity and connectivity in gene expression control. Nat Commun 16: 970.

Rhodes JDP, Feldmann A, Hernández-Rodríguez B, Díaz N, Brown JM, Fursova NA, Blackledge NP, Prathapan P, Dobrinic P, Huseyin MK, et al. 2020. Cohesin Disrupts Polycomb-Dependent Chromosome Interactions in Embryonic Stem Cells. Cell Rep 30: 820–835.e10.

Rickels R, Shilatifard A. 2018. Enhancer Logic and Mechanics in Development and Disease. Trends Cell Biol 28: 608–630.

Robinson JT, Thorvaldsdóttir H, Winckler W, Guttman M, Lander ES, Getz G, Mesirov JP. 2011. Integrative genomics viewer. Nat Biotechnol 29: 24–26.

Rostovskaya M. 2022. Maintenance of Human Naïve Pluripotent Stem Cells. Methods Mol Biol Clifton NJ 2416: 73–90.

Rostovskaya M, Stirparo GG, Smith A. 2019. Capacitation of human naïve pluripotent stem cells for multi-lineage differentiation. Dev Camb Engl 146: dev172916.

Rozowsky J, Gao J, Borsari B, Yang YT, Galeev T, Gürsoy G, Epstein CB, Xiong K, Xu J, Li T, et al. 2023. The EN-TEx resource of multi-tissue personal epigenomes & variant-impact models. Cell 186: 1493–1511.e40.

Schoenfelder S, Fraser P. 2019. Long-range enhancer–promoter contacts in gene expression control. Nat Rev Genet 20: 437–455.

Schoenfelder S, Furlan-Magaril M, Mifsud B, Tavares-Cadete F, Sugar R, Javierre B-M, Nagano T, Katsman Y, Sakthidevi M, Wingett SW, et al. 2015a. The pluripotent regulatory circuitry connecting promoters to their long-range interacting elements. Genome Res 25: 582–597.

Schoenfelder S, Javierre B-M, Furlan-Magaril M, Wingett SW, Fraser P. 2018. Promoter Capture Hi-C: High-resolution, Genome-wide Profiling of Promoter Interactions. J Vis Exp 57320.

Schoenfelder S, Sugar R, Dimond A, Javierre B-M, Armstrong H, Mifsud B, Dimitrova E, Matheson L, Tavares-Cadete F, Furlan-Magaril M, et al. 2015b. Polycomb repressive complex PRC1 spatially constrains the mouse embryonic stem cell genome. Nat Genet 47: 1179–1186.

Steindel M, Davis O, Neumann K, Pirvan L, Agsu G, Kranz A, Adhya D, Morf J, Yang S, Zhang Z, et al. 2025. A non-catalytic role for MLL2 in controlling chromatin organisation and mobility during the priming of pluripotent cells for differentiation. http://biorxiv.org/lookup/doi/10.1101/2025.02.21.639010.

Stock JK, Giadrossi S, Casanova M, Brookes E, Vidal M, Koseki H, Brockdorff N, Fisher AG, Pombo A. 2007. Ring1-mediated ubiquitination of H2A restrains poised RNA polymerase II at bivalent genes in mouse ES cells. Nat Cell Biol 9: 1428–1435.

Takashima Y, Guo G, Loos R, Nichols J, Ficz G, Krueger F, Oxley D, Santos F, Clarke J, Mansfield W, et al. 2014. Resetting transcription factor control circuitry toward ground-state pluripotency in human. Cell 158: 1254–1269.

Taylor T, Sikorska N, Shchuka VM, Chahar S, Ji C, Macpherson NN, Moorthy SD, de Kort MAC, Mullany S, Khader N, et al. 2022. Transcriptional regulation and chromatin architecture maintenance are decoupled functions at the Sox2 locus. Genes Dev 36: 699–717.

Theunissen TW, Powell BE, Wang H, Mitalipova M, Faddah DA, Reddy J, Fan ZP, Maetzel D, Ganz K, Shi L, et al. 2014. Systematic identification of culture conditions for induction and maintenance of naive human pluripotency. Cell Stem Cell 15: 471–487.

Thomson JA, Itskovitz-Eldor J, Shapiro SS, Waknitz MA, Swiergiel JJ, Marshall VS, Jones JM. 1998. Embryonic stem cell lines derived from human blastocysts. Science 282: 1145–1147.

Tian R, Abarientos A, Hong J, Hashemi SH, Yan R, Dräger N, Leng K, Nalls MA, Singleton AB, Xu K, et al. 2021. Genome-wide CRISPRi/a screens in human neurons link lysosomal failure to ferroptosis. Nat Neurosci 24: 1020–1034.

Vallot C, Hérault A, Boyle S, Bickmore WA, Radvanyi F. 2015. PRC2-independent chromatin compaction and transcriptional repression in cancer. Oncogene 34: 741–751.

Visel A, Minovitsky S, Dubchak I, Pennacchio LA. 2007. VISTA Enhancer Browser--a database of tissue-specific human enhancers. Nucleic Acids Res 35: D88–92.

Wang C, Lee J-E, Lai B, Macfarlan TS, Xu S, Zhuang L, Liu C, Peng W, Ge K. 2016. Enhancer priming by H3K4 methyltransferase MLL4 controls cell fate transition. Proc Natl Acad Sci 113: 11871–11876.

Wilkinson AL, Zorzan I, Rugg-Gunn PJ. 2023. Epigenetic regulation of early human embryo development. Cell Stem Cell 30: 1569–1584.

Wingett S, Ewels P, Furlan-Magaril M, Nagano T, Schoenfelder S, Fraser P, Andrews S. 2015. HiCUP: pipeline for mapping and processing Hi-C data. F1000Research 4: 1310.

Wu T, Hu E, Xu S, Chen M, Guo P, Dai Z, Feng T, Zhou L, Tang W, Zhan L, et al. 2021. clusterProfiler 4.0: A universal enrichment tool for interpreting omics data. The Innovation 2: 100141.

Xia W, Xie W. 2020. Rebooting the Epigenomes during Mammalian Early Embryogenesis. Stem Cell Rep 15: 1158–1175.

Xiang Y, Zhang Y, Xu Q, Zhou C, Liu B, Du Z, Zhang K, Zhang B, Wang X, Gayen S, et al. 2020. Epigenomic analysis of gastrulation identifies a unique chromatin state for primed pluripotency. Nat Genet 52: 95–105.

Xiong J, Zhu B. 2025. Epigenetic Preparation of Future Gene Induction Kinetics. Annu Rev Genet.

Xu B, On DM, Ma A, Parton T, Konze KD, Pattenden SG, Allison DF, Cai L, Rockowitz S, Liu S, et al. 2015. Selective inhibition of EZH2 and EZH1 enzymatic activity by a small molecule suppresses MLL-rearranged leukemia. Blood 125: 346–357.

Yu G. 2020. Gene Ontology Semantic Similarity Analysis Using GOSemSim. In Stem Cell Transcriptional Networks (ed. B.L. Kidder), Vol. 2117 of Methods in Molecular Biology, pp. 207–215, Springer US, New York, NY http://link.springer.com/10.1007/978-1-0716-0301-7_11.

Zaugg JB, Sahlén P, Andersson R, Alberich-Jorda M, de Laat W, Deplancke B, Ferrer J, Mandrup S, Natoli G, Plewczynski D, et al. 2022. Current challenges in understanding the role of enhancers in disease. Nat Struct Mol Biol 29: 1148–1158.

Zhang Y, Liu T, Meyer CA, Eeckhoute J, Johnson DS, Bernstein BE, Nusbaum C, Myers RM, Brown M, Li W, et al. 2008. Model-based analysis of ChIP-Seq (MACS). Genome Biol 9: R137.

Zheng H, Huang B, Zhang B, Xiang Y, Du Z, Xu Q, Li Y, Wang Q, Ma J, Peng X, et al. 2016. Resetting Epigenetic Memory by Reprogramming of Histone Modifications in Mammals. Mol Cell 63: 1066–1079.

Zijlmans DW, Talon I, Verhelst S, Bendall A, Van Nerum K, Javali A, Malcolm AA, van Knippenberg SSFA, Biggins L, To SK, et al. 2022. Integrated multi-omics reveal polycomb repressive complex 2 restricts human trophoblast induction. Nat Cell Biol 24: 858–871.

